# Shifts in protein aggregate stability define proteostasis decline in the aging human brain

**DOI:** 10.64898/2026.03.27.714902

**Authors:** Edward Anderton, Jordan B. Burton, Christina D. King, Anna C. Foulger, Dipa Bhaumik, Daria Timonina, Zachary Mayeri, Manish Chamoli, Julie K. Andersen, Birgit Schilling, Gordon J. Lithgow

## Abstract

Loss of proteostasis and the accumulation of insoluble protein aggregates are features of aging across model organisms and occur in all major age-related neurodegenerative diseases; yet how aggregation proceeds during normal human brain aging remains unknown. Using detergent-fractionation proteomics, we show that human brain aging does not involve uniform aggregate accumulation; rather, the insoluble proteome undergoes asymmetric remodeling beginning in midlife. Maximum-stability aggregates decline sharply in old age whereas intermediate-stability aggregates accumulate gradually before accelerating after age 80. Intermediate-stability aggregates are prone to liquid-liquid phase separation and are enriched with Alzheimer’s disease plaque and tangle constituents. Proteasome and cytosolic chaperone abundance predict individual differences in aggregate burden as strongly as age, offering supportive evidence, in humans, for therapies targeting these pathways. These findings establish aggregate remodeling as a feature of normal brain aging and position the accumulation of intermediate-stability aggregates as a molecular event on the path to neurodegenerative disease.

## Introduction

Loss of proteostasis is a conserved hallmark of aging^1^. When proteostasis mechanisms fail, cells and tissues accumulate misfolded proteins that form insoluble protein aggregates^2^. In animal models, aging is associated with the accumulation of detergent insoluble protein aggregates^3–7^. In humans, all major age-related neurodegenerative diseases involve the accumulation of detergent resistant, pathological aggregates^8^. Yet, whether insoluble proteins accumulate during normal human brain aging is unknown.

Insoluble protein aggregates exist on a continuum of stability defined by their biophysical and chemical properties^9–14^. Differential detergent fractionation has been used to characterize aggregated Tau, α-synuclein, prion protein, and TMEM106B in disease contexts, exploiting the principle that detergents of different strengths isolate aggregates based on their stability state^15–18^. N-lauroylsarcosine (sarkosyl) enriches aggregates maintained by intermediate-strength interactions, whereas sodium dodecyl sulfate (SDS), a stronger denaturant, leaves only the most resistant material insoluble ^14,19–22^. Sarkosyl-insoluble fractions enrich assemblies that retain biological activity such as seeding competence, which contributes to aggregate spread through tissues; SDS-resistant material, by contrast, is associated with highly stable, sequestered deposits ^20,23–28^. While the consequences of aggregate stability remain contested, soluble oligomers and intermediate-stability assemblies have been implicated as toxic species in neurodegeneration, in part because they retain seeding competence and can propagate through neural circuits^29,30^. Conversely, highly stable aggregates can limit toxicity by sequestering misfolded proteins from the functional pool^31,32^. Whether aging shifts the balance toward one stability state or another has not been examined in the human brain.

Protein biophysical analyses established that sequence and structural features increase aggregation propensity, including β-sheet-forming segments, hydrophobicity, intrinsic disorder, and motifs enabling multivalent interactions^33,34^. Classical amyloids, such as amyloid-β, aggregate through β-sheet-rich fibrillization, whereas other proteins, particularly RNA-binding proteins, assemble through liquid-liquid phase separation (LLPS) followed by maturation into solid aggregates^35–38^,. While these biophysical principles have been characterized extensively in disease contexts, whether they shape insoluble protein behavior during normal human brain aging is undetermined.

Here we asked whether normal human brain aging is characterized by the accumulation of insoluble protein aggregates. We applied parallel detergent fractionation with SDS and sarkosyl to post-mortem hippocampal tissue from disease-free individuals spanning the adult lifespan. Rather than uniform accumulation, we uncovered a qualitative reorganization of the insoluble proteome during aging, in which aggregates shift away from maximum-stability and towards intermediate-stability states. The proteins that accumulate possess a biophysical signature dominated by liquid-liquid phase separation propensity, rather than classical amyloid features, and are enriched among Alzheimer’s disease plaque and tangle constituents.

## Results

### Aging induces asymmetric remodeling of detergent-insoluble proteoforms in the female brain

We applied parallel detergent fractionation with SDS and sarkosyl to post-mortem hippocampal tissue from disease-free females aged 20-88 years (n = 52). Females were chosen due to their increased lifetime risk of neurodegeneration, in particular AD^39^. Subjects were excluded based on history of chronic age-related diseases, neuropsychiatric disorders, smoking, and substance abuse (see Methods for detailed inclusion-exclusion criteria). By extracting these fractions from the same tissue, we could directly compare how proteins change across stability states during brain aging. The insoluble proteome from each fraction, and lysate (soluble + insoluble), were quantified by mass spectrometry with data-independent acquisition (DIA). For differential abundance analysis, individuals were grouped: Young (20-35 yrs), Middle (45-55 yrs), Old (65-75 yrs), and Geriatric (80-88 yrs); post-mortem interval (PMI) did not differ across the groups (Kruskal-Wallis H = 1.33, p = 0.72) (Fig. 1A).

**Figure 1:**
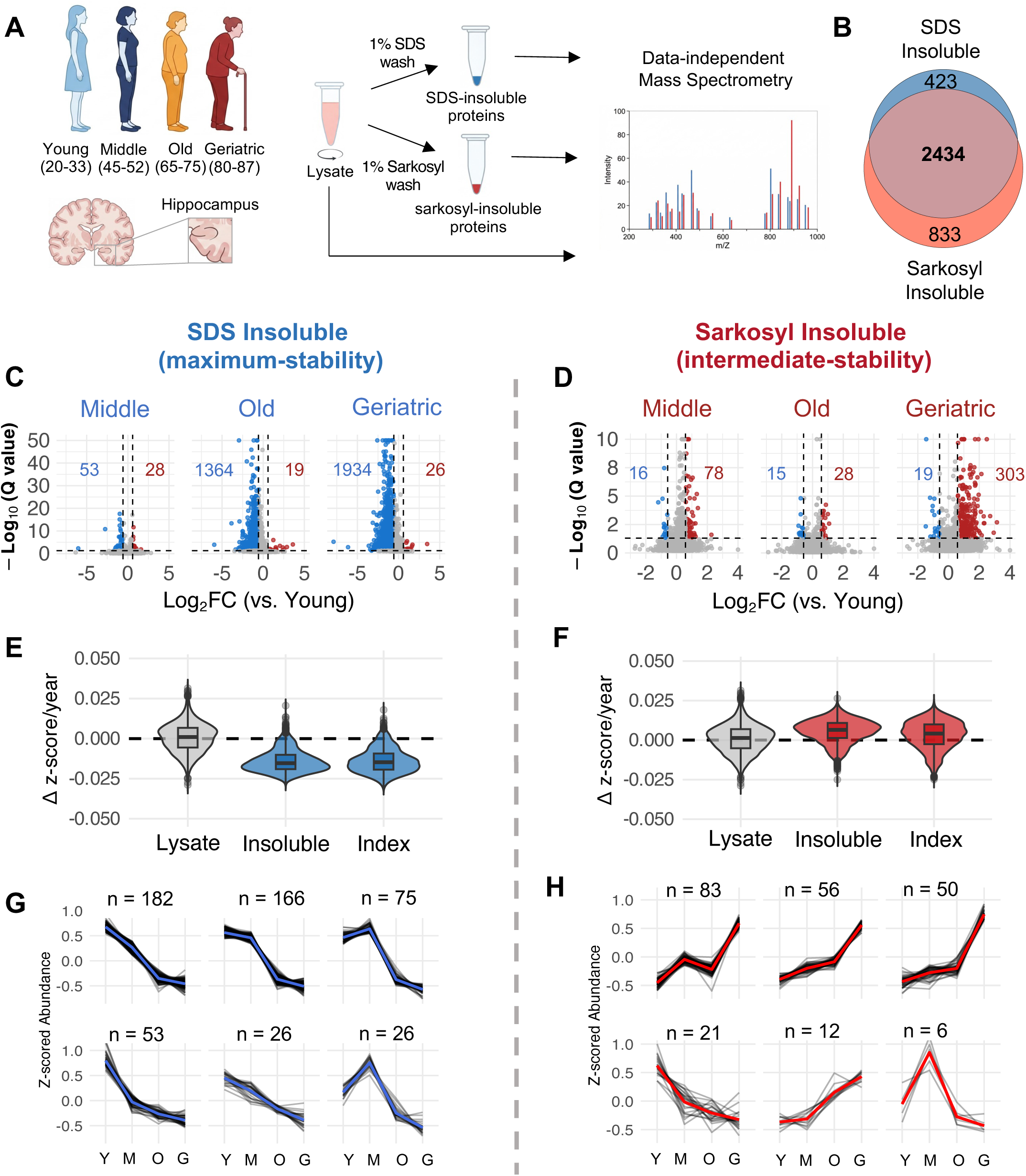
Brain aging causes proteome-wide, asymmetric remodeling of the insoluble aggregate proteome. **A**. Schematic representation of subject ages, insoluble protein fractionation with mass spec. workflow. **B.** Overlap of detected insoluble proteins in each fraction. **C.** Volcano plots of SDS insoluble protein abundance Log_2_(Fold Change) for young (n=12) vs. middle (n=12), vs. old (n=14), and vs. geriatric (n=14), Storey method with paired t-tests and p-values corrected for multiple testing using group-wise testing corrections, significance = |Log_2_(FC)| > 0.58, Q < 0.05). **D.** As in (C) with Sarkosyl insoluble protein abundance, n: young=9, middle=9, old=9, geriatric =11. **E.** Δ z score average change in abundance per year for every protein in lysate, SDS insoluble abundance and SDS insoluble index (insoluble / lysate). **F.** As in (E) with Sarkosyl insoluble proteome. **G.** Clustered trajectories of age-changing SDS insoluble proteins (k = 6). H. As in (G) with Sarkosyl insoluble proteome.

The two fractions captured largely overlapping proteins (overlap: 75-85%); however, they showed asymmetric age-related changes (Fig. 1B). The SDS-insoluble proteome decreased modestly from young to middle age but underwent a fraction-wide decrease in the transition to old age. In total, 82% of detected SDS-insoluble proteins decreased with age (n = 1934), representing general depletion of high-stability protein aggregates (Fig. 1C). TMEM106B, which forms bona fide amyloid fibrils in aged human brains^40^, was a notable exception, showing significant monotonic accumulation in the SDS-insoluble fraction across the lifespan. Conversely, sarkosyl-insoluble proteome changes were modest until the geriatric transition, after which approximately 10% of the proteome (n = 303 proteins) significantly increased (Fig. 1D). Total SDS-insoluble protein burden followed a log-linear decrease with age, whereas total sarkosyl-insoluble protein burden remained approximately constant (Supp. Fig 1D). Insoluble MAPT (Tau) showed depletion from SDS-insoluble aggregates alongside a trend towards accumulation in sarkosyl-insoluble aggregates (q=0.18) which showed high variance among geriatrics (Supp. Fig. 1E). Leave-one-out analysis and covariate-adjusted models confirmed that significant age-related changes were robust to individual subject exclusions, post-mortem interval, and biobank source; variation in effect magnitude was not associated with insoluble Amyloid Precursor Protein (APP) or Tau levels (see Methods: Robustness Estimations). Western blot analysis of a subset of samples confirmed the direction of age-related changes observed by mass spectrometry (Supp. Fig. 2A-D).

**Figure 2:**
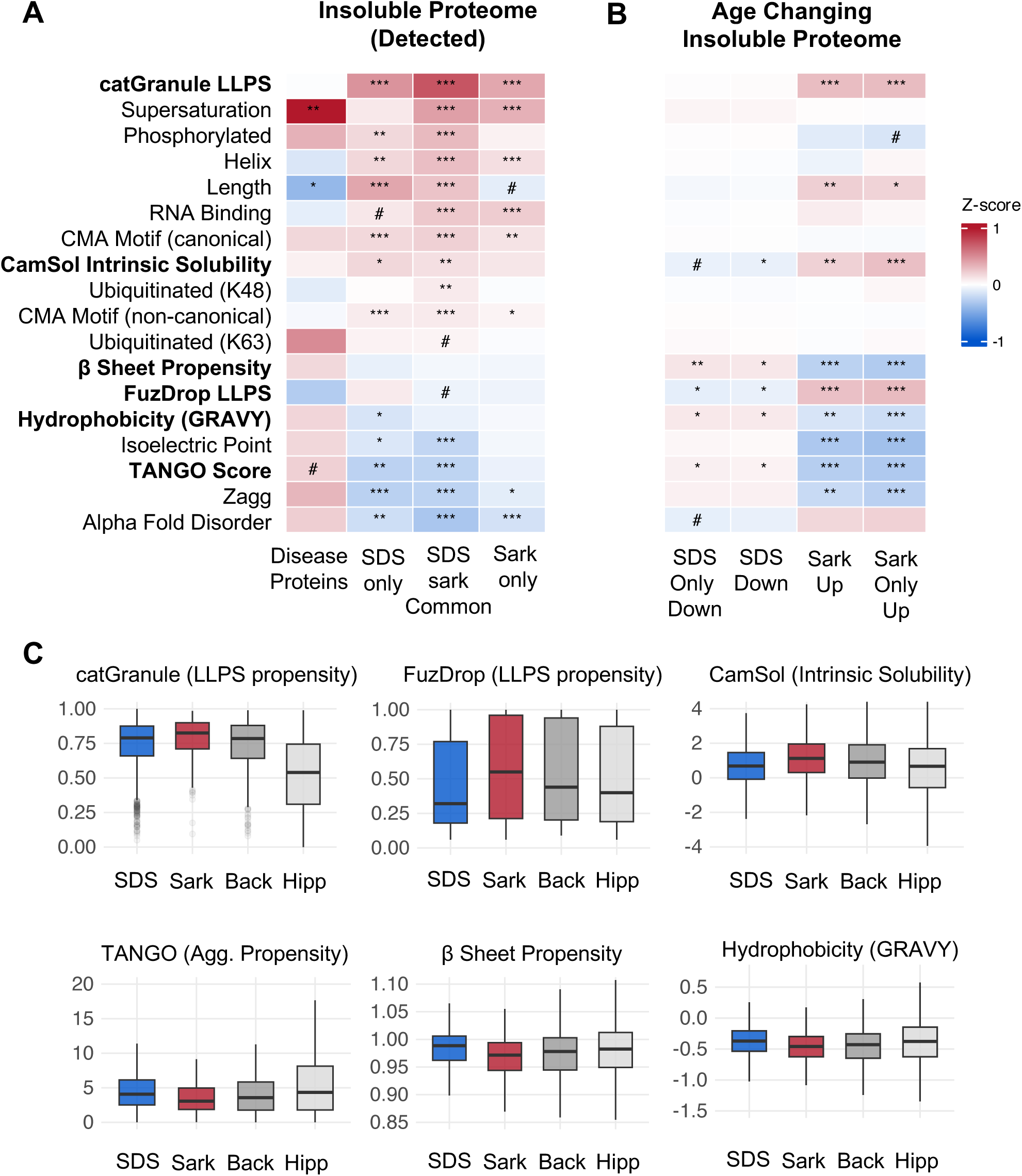
Liquid-liquid phase separation and intrinsic aggregation properties define aggregation state changes during brain aging. **A.** Heatmap showing enrichment of biophysical and functional features amongst insoluble proteins compared to the hippocampal proteome. Z-scores calculated as the difference between group and background central tendency (mean or median), normalized by background dispersion. Continuous variables: Welch’s t-test with Hedges’ g effect size (normal distribution) or Wilcoxon rank-sum test with Cliff’s delta (non-normal dist.), all use Benjamini-Hochberg FDR correction **B.** As in (A) with enrichment of age-changing proteins compared to insoluble detected proteins (non-changing). **C.** Boxplots for select features that show significant effects in the sarkosyl age-changing group. Extreme outlier proteins represented by grey points. Back = Insoluble detected proteins in SDS and Sarkosyl. Hipp – Hippocampal proteome. ***q < 0.001, **q < 0.01, *q < 0.05, #q <0.1.

Thousands of proteins were detected in both SDS- and sarkosyl-insoluble fractions across all age groups (SDS = 2857, Sark = 3267), spanning diverse biological processes (Fig. 1B, Supp. Fig. 1A). Synaptic proteins were the most over-represented cellular component in both fractions, followed by the endomembrane system and mitochondrial proteins (Supp. Fig. 1A). Both insoluble proteomes were enriched with proteins implicated across multiple proteinopathies, including AD, PD, Huntington’s disease, and prion disease (Supp. Fig. 1B). Proteins detected exclusively in the sarkosyl-insoluble fraction were enriched for mitochondrial ribosomes and ER proteins, whereas SDS-exclusive proteins showed no functional enrichment (Supp. Fig. 1C). Taken together, these data indicate that aging does not drive uniform aggregate accumulation; rather, it induces divergent, aggregation-state dependent changes across the proteome.

### Age-related insolubility changes are independent of expression

Protein solubility is tightly coupled to expression level, and aggregation can occur when abundance exceeds the solubility limit^41^. The observed age-related changes in insoluble protein could therefore reflect altered expression. To assess this, we calculated an insolubility index for each protein (insoluble/lysate) and computed age-dependent slopes for insoluble abundance, lysate abundance, and insolubility index separately (Fig. 1E&F). Lysate abundance was largely stable with age, and insoluble abundance slopes strongly predicted index slopes in both fractions (SDS: r = 0.83; sarkosyl: r = 0.76), establishing that changes in aggregate abundance rather than total expression underlie observed insolubility shifts (Fig. 1E&F). Given high concordance between insoluble abundance and insolubility index trajectories (SDS 95.9%, sarkosyl 80.0%), we chose to use insoluble abundance as the measure of aggregate load throughout.

The aging slope distributions showed a proteome-wide divergent pattern between the two solubility states: 95.2% of SDS-insoluble proteins declined with age, while 78.7% of sarkosyl-insoluble proteins increased (Fig. 1E&F). We hypothesized that this concurrent decline and accumulation could reflect redistribution between stability states. However, age-related changes were positively correlated across the proteome, and only a subset of SDS-declining proteins showed significant, concurrent sarkosyl accumulation, suggesting that protein is not redistributing from SDS-insoluble into sarkosyl-insoluble aggregates during aging (Supp. Fig. 3A).

**Figure 3:**
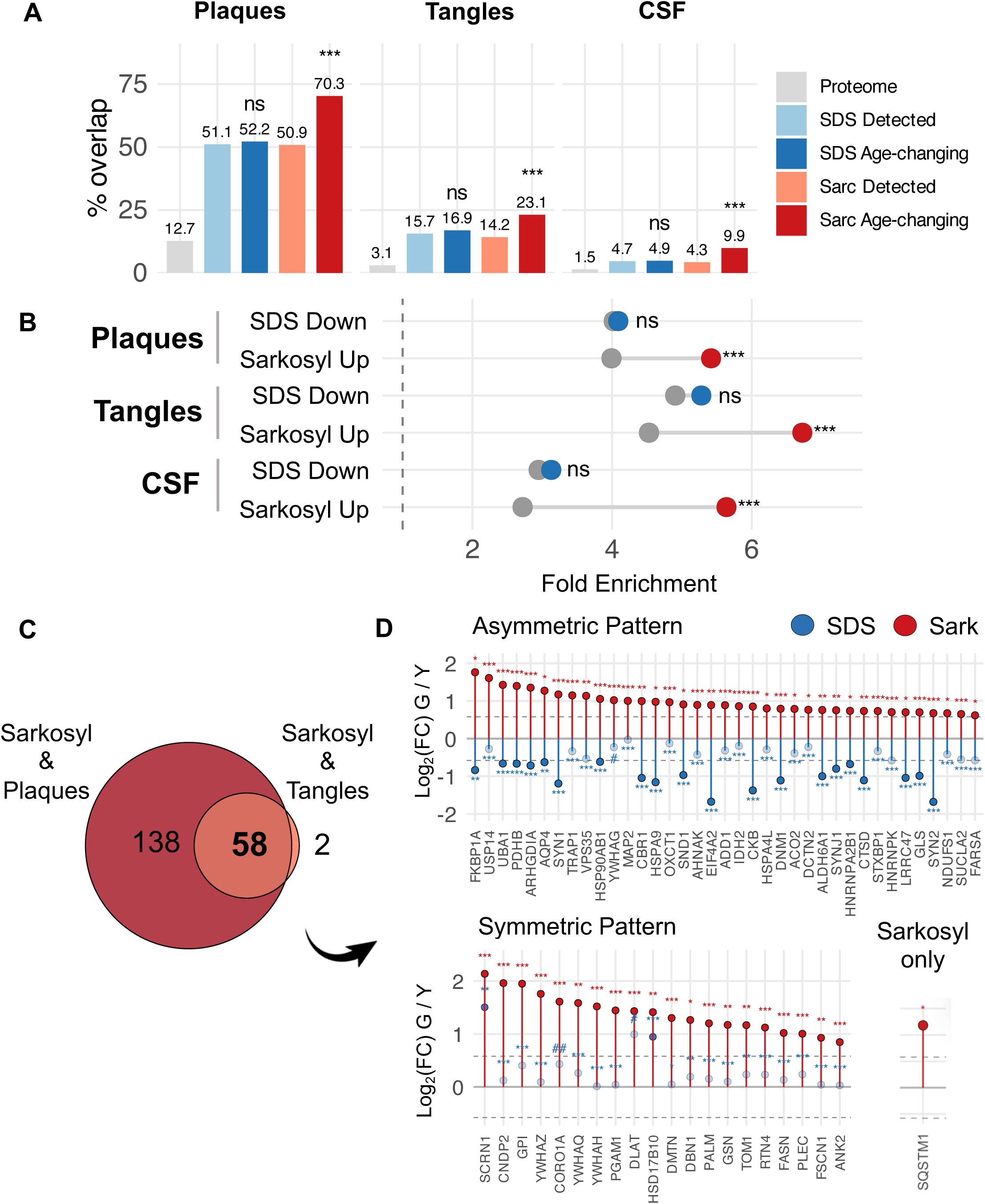
Proteins that accumulate in sarkosyl-insoluble aggregates are enriched with AD plaque, tangle and CSF proteins. **A**. Barplots showing percentage of insoluble proteome common with AD plaques, tangles, and CSF biomarkers. Proteome = Hippocampal proteome, Fischer’s exact test. **B.** Forest plot showing fold enrichment of insoluble proteome in AD plaques, tangles, and CSF biomarkers. Grey = insoluble detected, Blue = SDS age-changing proteins, Red = Sarkosyl age-changing proteins, Fisher’s exact test. **C**. Overlap of sarkosyl accumulators with plaques and tangles. **D** lolipop plots showing Log_2_(Fold Change) using geriatric vs young comparison for the 58 insoluble proteins common to sarkosyl accumulation, plaques and tangles. Blue = SDS, Red = Sarkosyl. Solid circle = significant, open circle = not significant. ***q < 0.001, **q < 0.01, *q < 0.05, #q <0.1, ##q <0.2,

The fraction-wide decline in SDS-insoluble abundance raised the question of whether tissue composition changes, such as synapse or neuronal loss, could account for the observed patterns. However, synaptic and postsynaptic density proteins showed stable lysate abundance with age while their SDS insolubility indices declined. Cell-type deconvolution confirmed that neuronal proportions were unchanged, ruling out synapse or neuronal loss as explanations (Supp. Fig. 4A, C, E). Interestingly, most synaptic proteins showed increases in sarkosyl insolubility index, consistent with a general increase within intermediate-stability aggregates during aging (Supp. Fig. 4B, D).

**Figure 4:**
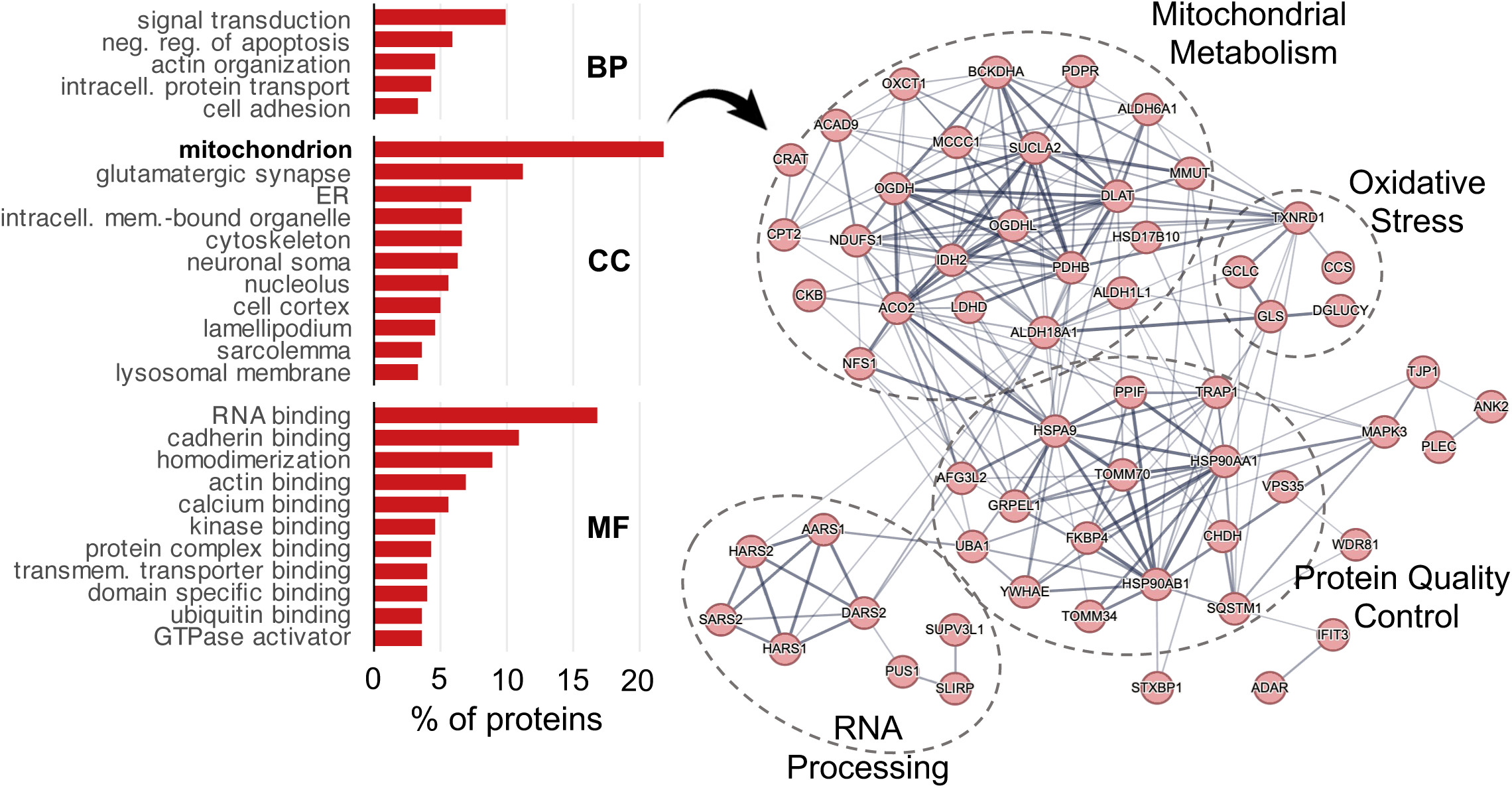
Mitochondrial and synaptic proteins accumulate in sarkosyl insoluble aggregates during brain aging. **A**. Barplots showing proportion of sarkosyl accumulating proteins across functional GO categories. **B**. STRING protein-protein interaction network diagram showing sarkosyl accumulating mitochondrial proteins.

### Midlife marks the onset of progressive aggregate remodeling

To identify the temporal pattern of insoluble protein changes across the lifespan, we fitted generalized additive models to each protein’s abundance as a function of age and clustered proteins with significant age effects by trajectory similarity (see Methods for filtering criteria). All age-changing SDS-insoluble proteins declined with age, with most following a biphasic pattern: slight declines from young to middle age and then steep declines in the middle-to-old transition (Fig. 1G). Conversely, most age-changing sarkosyl-insoluble proteins increased modestly from young through to old age before accelerating in the geriatric group (Fig. 1H). We applied a monotonic trend test to sarkosyl-insoluble proteins that accumulate during aging and found that 110 (36%) proteins reached significance after multiple testing correction (Mann-Kendall, q < 0.05; median p = 0.031), and all 303 proteins exhibited positive trend direction, suggesting that the accumulation of sarkosyl insoluble aggregates is a progressive process spanning decades.

### Selective accumulation of sarkosyl-insoluble aggregates is associated with liquid-liquid phase separation

We hypothesized that biophysical properties might distinguish proteins that decrease in maximum stability, SDS-insoluble aggregates or accumulate in intermediate stability, sarkosyl-insoluble aggregates during aging. To assess this, we applied a panel of established predictors of aggregation spanning intrinsic aggregation (TANGO, Zagg)^42,43^, predicted solubility (CamSol)^44^, supersaturation^45^, LLPS propensity (catGranule, FuzDrop)^46,47^, RNA-binding capacity^48^, biochemical and 3D structural properties^48^, AlphaFold predicted disorder^49^, post-translational modification sites^48^, and chaperone-mediated autophagy (CMA) degradation targeting motifs^50^(Fig. 2A). As a benchmark, we compared against 15 pro-aggregating, amyloids detectable in the hippocampus such as Amyloid β, Tau, and α-synuclein.

We first asked whether proteins vulnerable to insolubility, independent of age, share biophysical features with classical amyloids. For each protein group, we calculated feature enrichment relative to the hippocampal proteome (Figure 2A). Amyloids exhibited the expected aggregation signature: elevated TANGO scores, high β-sheet propensity, and low predicted solubility (Fig. 2A). The detergent-insoluble proteomes, by contrast, showed a largely opposite profile: lower aggregation propensity, reduced β-sheet content, and higher predicted solubility (Fig. 2A). Instead, insoluble proteins were characterized by elevated LLPS propensity (catGranule) and RNA-binding capacity (Fig. 2A). The connection with LLPS and RNA-binding capacity is particularly striking since many neurodegenerative disease proteins such as FUS, TDP-43, α-synuclein and HNRNPA1 associate with RNA-protein granules *in vivo*^38^. The single point of convergence between amyloids and the broader insoluble proteome was high supersaturation, indicating that expression near the solubility limit is a common feature of aggregation vulnerability (Fig. 2A).

We next asked whether age-changing proteins represent biophysically distinct subgroups. We performed biophysical enrichment of SDS-declining and sarkosyl-accumulating proteins, comparing each group against its detection background to account for baseline differences (Fig. 2B). The SDS-insoluble proteome was depleted for classical aggregation features relative to the hippocampal proteome (Fig. 2A). Within this depleted background, SDS-insoluble age-declining proteins showed modest relative enrichment for TANGO scores, beta-sheet propensity, and hydrophobicity, indicating that the proteins lost during aging are the most aggregation-prone members of their fraction (Fig. 2B). By contrast, the sarkosyl-insoluble age-accumulating proteins were depleted of classical aggregation features but enriched for LLPS propensity (catGranule and FuzDrop), consistent with condensate-mediated aggregation (Fig. 2B). While both LLPS predictors were enriched among sarkosyl-accumulating proteins, catGranule and FuzDrop scores were only modestly correlated (r=0.29) which is consistent with their distinct training datasets. FuzDrop predicts based on entropy-driven disordered interactions, whereas catGranule is a broader phase separation predictor incorporating RNA-binding and structural features. Among sarkosyl-accumulating proteins, 79% scored in the top third of proteins for at least one LLPS predictor, partitioning into two approximately equal populations: proteins scoring high on catGranule but not FuzDrop (36%), and proteins scoring high on both predictors (37%). These populations, therefore, represent distinct routes to LLPS-driven aggregation, one mediated by entropy-based disordered interactions captured by both predictors and one driven by features such as multivalent interactions and RNA-binding capacity that FuzDrop does weight. Because LLPS predictors correlate with sequence length, we used multivariate logistic regression to confirm that both catGRANULE and FuzDrop independently distinguished accumulators after controlling for sequence length, abundance, and peptide counts (see Methods). Taken together, these data demonstrate that age-related sarkosyl accumulation is more likely to be driven by LLPS propensity than classical amyloid features, with two biophysically distinct subpopulations converging on condensate-mediated aggregation.

### Proteins that accumulate in sarkosyl-insoluble aggregates during aging are enriched in AD plaques and tangles

Sarkosyl is widely used to study pro-aggregating proteins in neurodegenerative disease, and several studies have reported alterations to the sarkosyl-insoluble proteome in AD brains^51–55^. The accumulation of sarkosyl-insoluble protein in the geriatric group, when AD risk increases sharply, led us to question whether this material resembles disease-associated aggregates. We compared our data against the NeuroPro database, which compiles AD proteomic studies including laser microdissection of amyloid plaques and neurofibrillary tangles^56^.

We first determined fold enrichment of proteins vulnerable to insolubility, irrespective of age. Both insoluble proteomes showed enrichment for plaque-associated proteins compared to the hippocampal proteome, with roughly half of insoluble-detected proteins appearing in the plaqueome; tangle enrichment followed a similar pattern with lower absolute representation (Fig. 3A&B, Supp. Fig. 5A&B). When comparing age-changing proteins in each insoluble fraction a clear dichotomy emerged. Proteins declining in the SDS-resistant, maximally stable fraction during aging showed no additional enrichment for plaques or tangles beyond baseline (Fig. 3A&B, Supp. Fig. 5C). Proteins accumulating in the intermediate-stability sarkosyl-insoluble pool, by contrast, showed pronounced enrichment for both AD plaque and tangle constituents (Fig. 3A&B, Supp. Fig. 5D). This result was remarkable since it suggests that aggregate accumulation during disease-free aging overlap significantly with AD.

**Figure 5.**
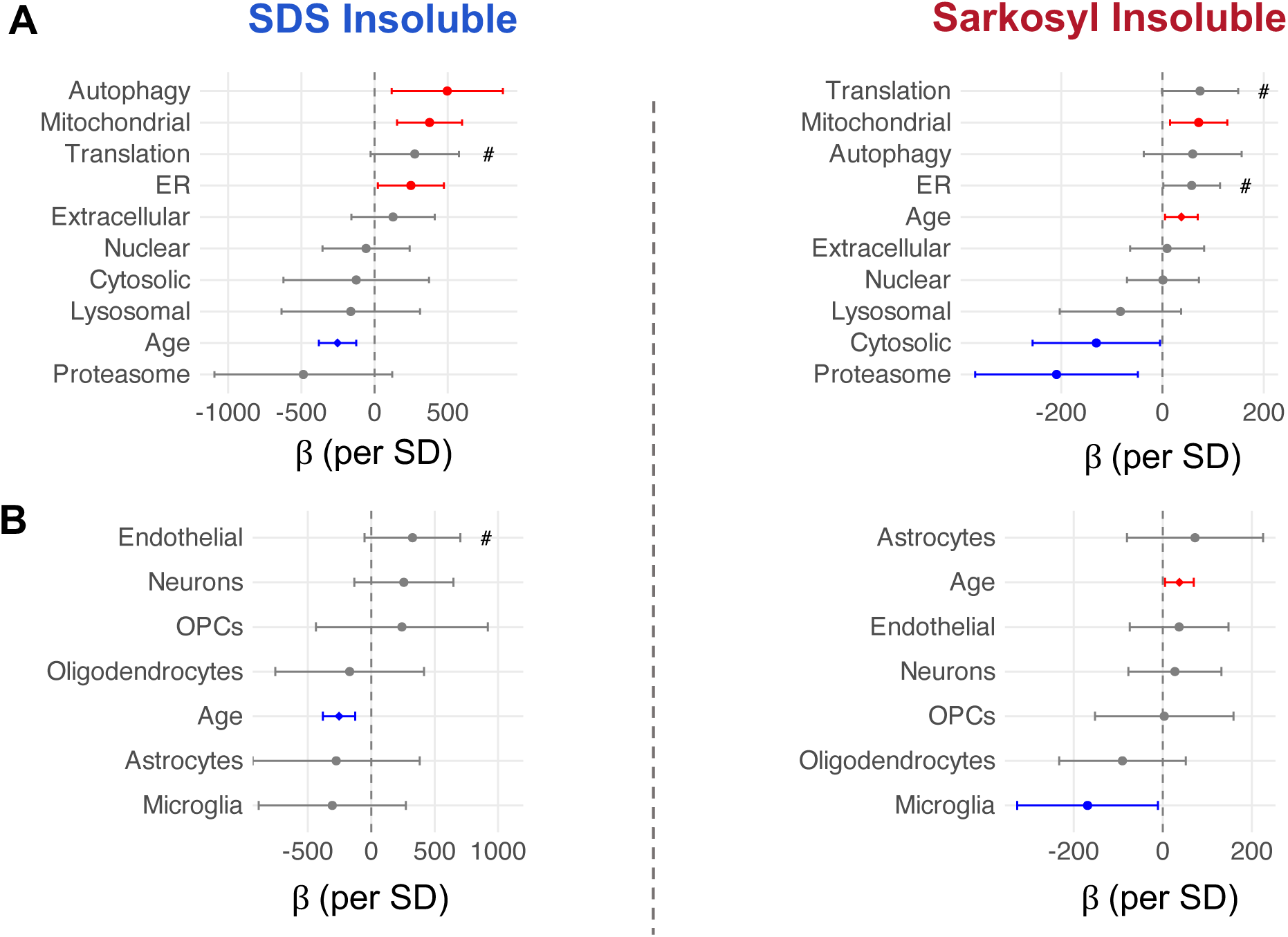
Proteostasis network capacity and cell type composition predict individual insoluble protein burden. **A** Forest plots showing the association between lysate-level proteostasis branch signature scores and the burden of age-changing insoluble proteins Left = SDS, Right = Sarkosyl. Points show the beta coefficient +/- 95% confidence interval from a linear model adjusting for age (insol. burden ∼ age + predictor), Red = significant positive predictor, Blue = significant negative predictor. **B.** As in (A), age-adjusted linear model but using cell type signature scores and insoluble burden. OPC = Oligodendrocyte Precursor Cells, #p <0.1.

We next asked whether sarkosyl-accumulating proteins have been identified as AD CSF biomarker candidates, using a meta-analysis of putative biomarkers identified across at least three independent discovery cohorts^57^. CSF biomarkers were enriched at baseline in both detected insoluble proteomes; however, only proteins that accumulate in the sarkosyl-insoluble fraction during aging showed significant enrichment above baseline, extending the disease relevance of age-related sarkosyl accumulation beyond brain tissue to peripheral biomarker signatures (Fig. 3A&B).

Fifty-eight proteins that increased in the sarkosyl-insoluble fraction with age were independently detected in both amyloid plaque and neurofibrillary tangle proteomes. Most of these (37/58, 64%) showed concurrent depletion from the SDS-insoluble fraction, consistent with loss from maximally stable aggregates alongside accumulation in intermediate-stability states (Fig. 3C). The remainder were either stable in the SDS-insoluble fraction or showed a trend toward higher SDS-insoluble levels during aging. These 58 proteins encompassed synaptic components spanning presynaptic machinery, postsynaptic cytoskeletal markers, and four major brain 14-3-3 isoforms; proteostasis machinery including SQSTM1(p62), VPS35, USP14, and cytosolic and mitochondrial chaperones including HSP90AB1, HSPA4L, HSPA9, and TRAP1; mitochondrial enzymes of the TCA cycle, pyruvate dehydrogenase complex, and oxidative phosphorylation; and RNA-binding proteins implicated in aberrant granule formation (HNRNPA2B1, HNRNPK)^58,59^ (Fig. 3C).

### Sarkosyl-insoluble aggregates span synaptic, mitochondrial, and proteostasis machinery in the aging female hippocampus

Because aggregation depletes functional protein, we asked which biological functions are represented among sarkosyl-insoluble accumulators. Functional annotation of all 303 accumulating proteins revealed nine broad categories (Table 1).

**Table 1.**
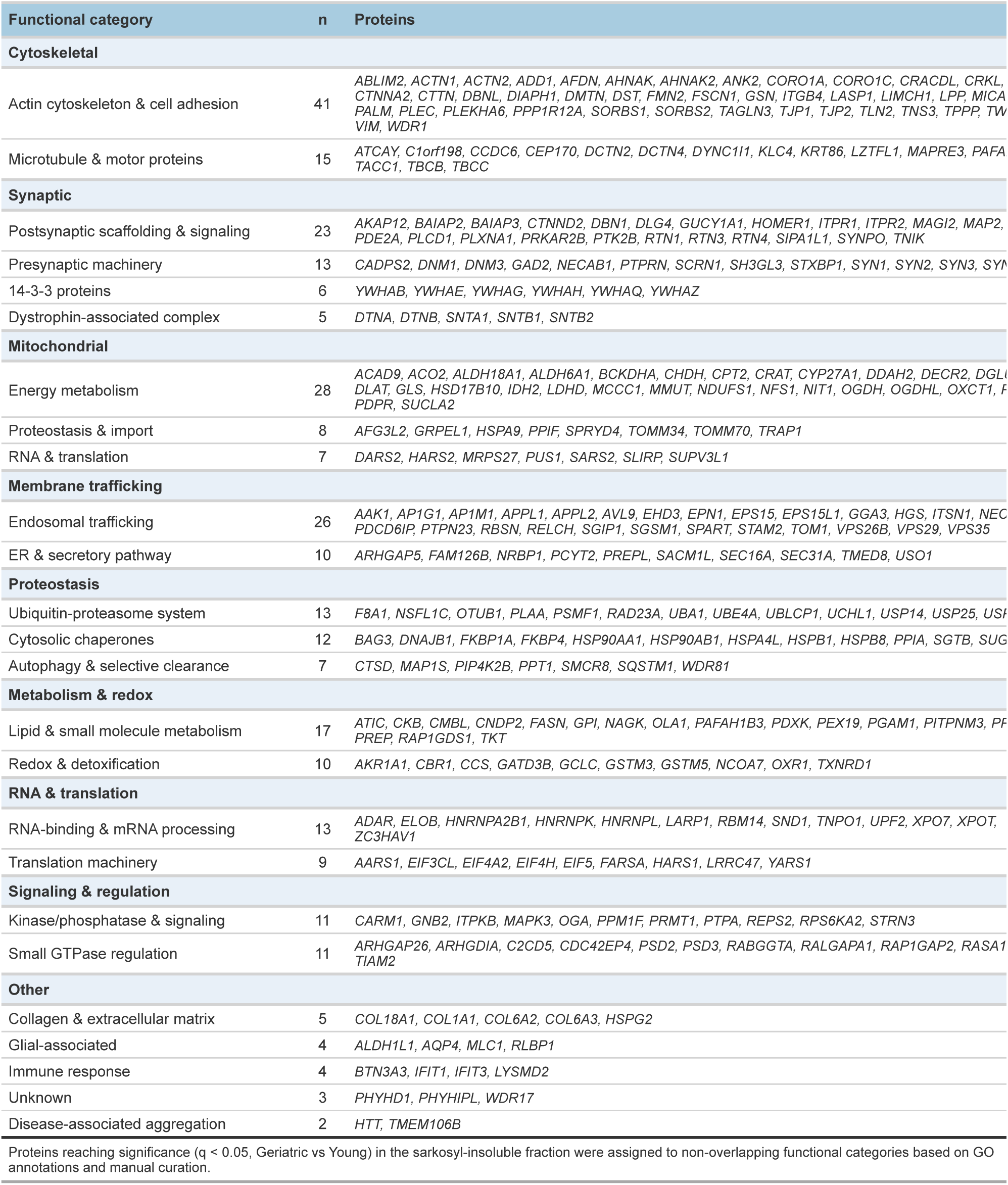
Functional classification of sarkosyl-insoluble accumulators (n = 303)

Cytoskeletal and synaptic proteins were the largest functional group, with actin cytoskeleton and cell adhesion components (n = 41), postsynaptic scaffolding and signaling proteins (n = 23), presynaptic machinery (n = 13), microtubule and motor proteins (n = 15), all six brain 14-3-3 isoforms, and the complete neuronal dystrophin-associated complex (Table 1).

Mitochondrial proteins were the second largest category (n = 43), including energy metabolism enzymes (n = 28), proteostasis and import machinery (n = 8), and mitochondrial RNA and translation factors (n = 7) (Table 1). Age-accumulating mitochondrial proteins were predominantly matrix enzymes, particularly TCA cycle components and quality control chaperones HSPA9 (mtHsp70) and TRAP1 (mtHsp90); membrane-integrated complexes of the electron transport chain, ATP synthase, and TIM translocase were, by contrast, constitutively detergent-resistant at all ages. The co-aggregation of TOMM70, which coordinates substrate recognition at the outer membrane import channel, alongside the matrix chaperones and enzymes it helps deliver, raises the possibility that protein import represents a bottleneck at which misfolded cargo accumulates (Table 1).

The remaining categories included membrane trafficking proteins (n = 36), among them VPS35, the retromer cargo-recognition subunit linked to familial Parkinson’s disease^60^; proteostasis machinery (n = 32) including ubiquitin-proteasome components, cytosolic chaperones, and selective autophagy receptors such as SQSTM1/p62; and RNA-binding and translation proteins (n = 22) (Table 1). Consistent with the LLPS and RNA-binding signature identified in the biophysical analysis, this last group included mRNA processing factors, translation initiation components, and aminoacyl-tRNA synthetases from both cytoplasmic and mitochondrial compartments, suggesting that mitochondrial translational capacity is itself vulnerable to age-related aggregation (Table 1). Taken together, the proteins accumulating in sarkosyl-insoluble aggregates during aging represent core machinery required for synaptic function, suggesting that proteostasis decline during aging may erode synaptic activity independently of synapse loss.

### Proteostasis machinery is enriched in insoluble fractions

The accumulation of mitochondrial quality control machinery in sarkosyl-insoluble aggregates during aging led us to question whether proteostasis machineries were enriched in either the SDS- or sarkosyl-insoluble protein aggregates. Their presence could provide clues as to which machineries handle the different aggregate species, and which could be vulnerable to co-aggregation during aging. To test this, we calculated composite abundance signatures for nine proteostasis network (PN) branches using the proteostasis network consortium database annotations^61,62^. Autophagy machinery and cytosolic chaperones were significantly enriched in both insoluble fractions across all ages (Supp. Fig. 6A). Most PNs were present in aggregates but remained stable across the lifespan (Supp. Fig. 6A). Autophagy and cytosolic chaperone proteins were enriched in both insoluble fractions irrespective of age, indicating that these machineries constitutively co-fractionate with aggregates (Supp. Fig. 6A). The two fractions diverged, however, with respect to age-related changes: autophagy components were enriched among SDS-declining proteins but depleted from sarkosyl-insoluble accumulators (Supp. Fig. 6A). This asymmetry is consistent with a model in which autophagy actively clears SDS-resistant aggregates and declines alongside them, while sarkosyl-insoluble material accumulates through a pathway that autophagy does not effectively engage. Mitochondrial protein homeostasis showed a trend towards enrichment in the sarkosyl age-changing insoluble proteome consistent with the enrichment of mitochondrial proteins in that fraction (Fig. 4, Supp. Fig. 6A).

We applied the same approach to CNS cell types using published composite cell type marker data^63^. Neuronal proteins were enriched in both insoluble fractions, microglial proteins were depleted across both, while astrocyte markers trended toward enrichment among sarkosyl-insoluble accumulators (Supp. Fig. 6B). This pattern indicates that the insoluble proteome is neuronally dominated at all ages but that astrocytic proteins may be selectively vulnerable to age-related accumulation while microglial proteins are excluded.

### Proteostasis network capacity predicts hippocampal aggregate burden independently of chronological age

While aging has a significant effect on insoluble protein levels, aggregate burden was highly heterogeneous even among age-matched individuals (Supp. Fig. 1D), suggesting that other factors contribute to aggregate load. We hypothesized that PN capacity could explain inter-individual variation in aggregation. To quantify this, we tested each PN signature score against insoluble burden in a linear model that also included age; coefficients therefore reflected proteostasis effects on aggregate accumulation independent of chronological age.

Two pathways showed significant protective associations with sarkosyl-insoluble accumulation: proteasome capacity yielded the largest individual effect size, followed by cytosolic chaperones (Fig. 5A, β per SD <0). We applied the same approach to cell-type signatures and found that microglia showed a protective association with sarkosyl-insoluble burden consistent with literature implicating microglia in aggregate clearance in the CNS^64^ (Fig. 5B).

Mitochondrial proteostasis capacity, by contrast, showed a positive association with insoluble burden in both fractions (Fig. 5A, β per SD >0). We ruled out mitochondrial mass as a predictor using total mitochondrial protein signal as a proxy, indicating that the association is specific to proteostasis capacity rather than organelle abundance. Autophagy and ER proteostasis showed significant positive associations with SDS-insoluble burden; ER also showed a trend for sarkosyl and translation showed positive trends in both fractions (Fig. 5A). The positive direction of these biosynthesis and import-associated pathways contrasted with the protective direction of clearance mechanisms, suggesting that protein production and import track with aggregate load while degradation capacity limits it.

The translation category encompassed hundreds of functionally heterogeneous proteins and so we decomposed it into bona fide translation machinery and co-translational proteostasis proteins (Supp. Fig. 6C). Consistent with the broader pattern, abundance of mitochondrial and cytosolic ribosomes was positively associated with insoluble protein levels across both fractions (Supp. Fig. 6C). Mitochondrial ribosome biogenesis was specifically associated with sarkosyl-insoluble accumulation, and mitochondrial protein import machinery was positively associated with insoluble protein across both fractions (Supp. Fig. 6C). Conversely, cytosolic elongation and termination factors and ribosome-associated nascent chain chaperones (NAC/RAC complex) both displayed protective associations, implicating co-translational proteostasis as a determinant of aggregate load (Supp. Fig. 6C). Taken together, these data indicate that aggregate burden in human brain reflects a balance between protein production and clearance capacity, with proteasome, cytosolic chaperones, and co-translational quality control limiting accumulation while biosynthetic and import pathways track with load.

## Discussion

Our proteome-wide analysis reveals that human brain aging does not drive uniform aggregate accumulation; rather, the insoluble proteome undergoes qualitative remodeling, with maximally stable SDS-resistant aggregates declining while intermediate-stability sarkosyl-insoluble proteins accumulate progressively before accelerating after age 80. Sarkosyl-accumulating proteins possess a biophysical signature centered on LLPS propensity rather than classical amyloid features; however, accumulating proteins show strong enrichment among AD plaque and tangle constituents. Aggregate burden varied substantially among age-matched individuals, with proteostasis network capacity explaining as much variance as chronological age, implying that aggregate accumulation is not an inevitable feature of aging. Because the proteins suppressed by high proteostasis capacity are the same proteins enriched in AD pathology, individual differences in clearance capacity may influence accumulation of disease-relevant aggregates during normal aging.

The temporal separation of these changes implies that the decrease in abundance of proteins in SDS insoluble aggregates and accumulation in sarkosyl insoluble aggregates are independent remodeling events rather than sequential phases of a single process. SDS-insoluble aggregates decrease most dramatically during the middle-to-old transition, whereas sarkosyl-insoluble protein accumulation remains gradual until the geriatric transition, when it accelerates markedly. The net effect is a shift in the aggregate landscape away from sequestered, maximally stable deposits and toward labile, intermediate-stability material associated with seeding competence in disease contexts. The surge in sarkosyl-insoluble aggregates at the geriatric transition may represent a threshold phenomenon in which clearance mechanisms become overwhelmed, though our cross-sectional design cannot test this directly. AD-associated pathology develops biochemically for decades before cognitive symptoms manifest^65^, and it is the AD-enriched sarkosyl-insoluble fraction that accelerates in the ninth decade when AD incidence rises sharply.

The proteome-wide decline of SDS-insoluble aggregates was unexpected given that aging in *C. elegans* and mice is associated with SDS-insoluble accumulation^4,66,67^. One possible interpretation is that maximally stable aggregates represent protective detention structures like aggresomes. Indeed, sequestration of soluble Aβ oligomers and HTT peptides into insoluble deposits reduces toxicity^32,68^. However, the SDS-insoluble proteome lacked classical aggresome markers: HDAC6^69^ was not detected, and SQSTM1/p62^70^ and BAG3^71^ appeared exclusively in sarkosyl-insoluble material. An alternative interpretation is that the SDS-insoluble fraction we measure represents a snapshot of material passing through autophagy and the age-related decline in SDS insoluble material reflects reduced flux. This is supported by the fact that autophagy components are enriched among age-declining SDS-insoluble proteins, and autophagy pathway expression tracks positively with SDS-insoluble burden. These models are not mutually exclusive because sequestered aggregates should be targeted to autophagy. Whether SDS-insoluble material in the human brain serves a protective function, represents a clearance intermediate, or both, remains to be determined.

Sarkosyl-accumulating proteins possess a biophysical signature distinct from classical amyloids. Rather than elevated beta-sheet propensity and hydrophobicity, they are characterized by greater sequence length, high LLPS propensity across two orthogonal predictors, and higher predicted solubility. LLPS proceeds through weak multivalent interactions largely independent of surface hydrophobicity, and sequence length correlates with multivalency^72^. The LLPS predictor profiles for accumulating proteins partition into two populations, suggesting distinct biophysical routes converging on condensate formation. RNA-binding proteins constitute the largest functional category among accumulators; however, RNA-binding capacity is not enriched above the sarkosyl-detected background, suggesting that LLPS-driven aggregation must proceed through broader mechanisms not restricted to canonical RNA-protein interactions. The enrichment of LLPS-prone proteins among AD plaque and tangle constituents suggests that phase separation may represent a pathway to pathological aggregation operating alongside classical amyloid formation. It has been proposed that condensates can mature into solid aggregates over time^38^, providing a plausible route from phase separation to detergent-insoluble material during aging, although the mechanism for solidification is unknown.

Aggregate burden was highly heterogeneous among age-matched individuals, and proteostasis network machinery abundance explained substantial inter-individual variance. Proteasome and cytosolic chaperone abundance showed negative, protective associations with sarkosyl-insoluble burden, whereas biosynthetic and import pathways showed positive associations with aggregate load. When combined in a linear model proteostasis predictors tripled the variance explained by age alone, predicting almost 40% of total variance in insoluble burden. Contrary to expectation, autophagy network abundance was not protective against sarkosyl-insoluble accumulation. Moreover, autophagy components were depleted among sarkosyl accumulators but enriched among age-declining SDS-insoluble proteins, suggesting that autophagy preferentially interacts with SDS-insoluble material. The protective association of proteasome capacity aligns well with model organism studies which implicate UPS decline as a driver of age-related aggregation^73^. Given that the proteins suppressed by high proteasome capacity are strongly enriched among AD plaque and tangle constituents, these data provide human-level evidence that UPS capacity may influence disease-relevant aggregate accumulation.

Mitochondrial proteostasis is emerging as a conserved age-related vulnerability, with mitochondrial protein aggregation now documented across *C. elegans*, mouse, and now human brain^3–5,67,74^. Mitochondrial proteins were prominent were prominent in the insoluble proteome but only matrix enzymes and quality control machinery showed age-related accumulation; membrane-integrated ETC complexes, ATP synthase, and TIM translocase were insoluble at all ages. One possible explanation for mitochondrial protein vulnerability is precursor over-accumulation stress (mPOS): when mitochondrial import capacity declines due to mitochondrial dysfunction, nuclear-encoded mitochondrial proteins accumulate in the cytosol, where they are prone to aggregation^75^. In this context, the link we observe between mitoribosome biogenesis factor abundance and overall sarkosyl-insoluble burden makes sense since biogenesis demand necessarily correlates with the pool of import-dependent precursors vulnerable accumulation in the cytosol. This model implies a feedforward loop in which mitochondrial dysfunction and mitochondrial protein aggregation reinforce one another, whether aggregation occurs in the cytosol following failed import or within the matrix itself.

This study has limitations worth acknowledging. Our cohort consisted exclusively of females, selected for their higher lifetime AD risk^76^ and whether males show equivalent aggregation patterns should be addressed in future work. While subjects were screened for neurodegenerative disease by neuropathological examination, we cannot exclude subcellular pathology, and variance in insoluble amyloid and tau among the oldest old may partly reflect preclinical heterogeneity. All subjects were cognitively normal at death, but we lack longitudinal cognitive data and cannot assess the functional relevance of insoluble protein changes. Our biophysical analysis was based on sequence-level predictors; we did not perform ultrastructural or material-state validation, so assignments of LLPS behavior remain to be demonstrated experimentally. Future work should prioritize biochemical characterization, expansion to male cohorts, and validation in murine models.

This study establishes that disease-free human brain aging involves qualitative remodeling of the insoluble proteome rather than uniform aggregate accumulation, with maximally stable and intermediate-stability aggregate pools showing independent and opposing trajectories. The proteins that accumulate in intermediate-stability aggregates during normal aging are enriched in AD plaques and tangles, and their accumulation is constrained by proteostasis capacity, particularly proteasome and cytosolic chaperone abundance. These observations provide evidence, in humans, that loss of proteostasis links normal aging to neurodegenerative disease and identify potential targets for intervention before clinical symptoms emerge.

To facilitate community re-use and allow readers to explore the datasets at the level of individual proteins, we provide the Human Hippocampus Insoluble Proteome Explorer, an interactive Shiny application for browsing age trajectories, biophysical features, and AD pathology overlap across the SDS-insoluble, sarkosyl-insoluble, and lysate proteomes https://jzezxu-edward-anderton.shinyapps.io/human-hippocampus-insoluble-proteome-explorer/.

## Methods

All statistical tests were performed using R base package unless otherwise stated.

### Human brain tissue sourcing

Human brain samples were sourced from the NIH Neurobiobank consortium, which encompasses several biobanks across the contiguous United States. Biobanks follow the same standardized sample processing and handling procedures.

### Subject inclusion / exclusion criteria

Frozen post-mortem human coronal tissue sections encompassing the entire hippocampal formation from the level of the lateral geniculate nucleus were obtained from 52 female donors aged 20-88 years. PMI did not differ across age groups (Young, median 19.0 h, IQR 16.3 to 23.3; Middle, 17.7 h, IQR 15.8 to 20.8; Old, 19.0 h, IQR 15.0 to 20.8; Geriatric, 21.5 h, IQR 13.2 to 24.9; Kruskal-Wallis H = 1.33, p = 0.72). Females were chosen due to their increased lifetime risk of neurodegeneration, in particular AD^39^. All hippocampus samples were deemed disease free by expert neuropathologists. To isolate the effects of normal aging on protein aggregation from confounding disease processes, we applied stringent exclusion criteria. Donors with a clinical history of neurodegenerative disease (Alzheimer’s disease, Parkinson’s disease, dementia), cerebrovascular events (stroke, transient ischaemic attack), psychiatric disorders (depression, anxiety, bipolar disorder, schizophrenia), substance abuse, or smoking were excluded. We further excluded donors with systemic inflammatory or autoimmune conditions (rheumatoid arthritis, autoimmune disorders, hepatitis A/B/C), organ or bone marrow transplant recipients, and those with major organ disease including cardiovascular disease (myocardial infarction, coronary artery disease, congestive heart failure, cardiomyopathy), chronic pulmonary disease (emphysema, COPD, pulmonary fibrosis, sleep apnoea), gastrointestinal disease (coeliac disease, Crohn’s disease), hepatic disease (cirrhosis, NAFLD/NASH), chronic kidney disease, diabetes, and macular degeneration. A subset of common conditions was included: asthma, osteoarthritis, fibromyalgia, endometriosis, IBS, GERD, Barrett’s oesophagus, glaucoma, cataracts, venous thromboembolism, kidney stones, and non-CNS cancers without brain metastases. Hypertension and hypercholesterolaemia were included due to the very high overall prevalence amongst the elderly^77^.

### Brain tissue homogenization

Homogenization and processing of the lysates was completed following universal precautions in a BSL2 certified tissue culture hood. A segment of hippocampal tissue encompassing all regions of the hippocampal formation was submerged in 150 µL of lysis buffer containing protease, phosphatase and deacylase inhibitors (20 mM Tris base, pH 7.4, 100 mM NaCl, 1 mM MgCl₂, 30 mM NAD, 5 mM Trichostatin A) per mg of frozen tissue in a pre-cooled 7 mL Dounce homogenizer, on ice. To homogenize the tissue, 30 strokes were applied with pestle A, followed by 30 strokes with pestle B. Homogenate was removed into a pre-cooled 15 mL conical tube and sonicated in a 4°C water bath using the Bioruptor for 5 cycles of 30 seconds on and 60 seconds off at medium intensity. To remove cell debris, the lysate was centrifuged in a pre-cooled, 4°C swinging bucket centrifuge at 3000×g for 4 minutes. The supernatant (lysate) was removed and stored at -80°C until subsequent processing. The pellet was discarded.

### Insoluble protein isolation

Samples were processed in batches of eight and, to minimize batch processing effects, an equal number of subjects from each of the pre-defined age groupings were processed in each batch. Subjects were selected randomly for each batch. The lysate protein concentration was quantified by BCA according to manufacturer’s instructions. To normalize input for insoluble protein isolation, an aliquot of lysate equivalent to 2 mg of total protein was added to a sterile, ice-cold Eppendorf tube and the volume was equalized across all samples by addition of ice-cold lysis buffer containing protease, phosphatase inhibitor and deacylase inhibitor (20 mM Tris base, pH 7.4, 100 mM NaCl, 1 mM MgCl2, 30mM NAD, 5mM Trichostatin A), typically to 1.2 mL. The protein solution was centrifuged in a 4°C pre-cooled benchtop centrifuge at 20,000×g for 15 minutes to pellet aqueous-insoluble proteins. The supernatant was collected as the aqueous-soluble fraction and stored at -80°C. The insoluble pellet was incubated in 1 mL of 1% (w/v) SDS or 1% (w/v) sarkosyl lysis buffer containing protease and phosphatase inhibitor ((20 mM Tris base, pH 7.4, 100 mM NaCl, 1 mM MgCl2, 30mM NAD, 5mM Trichostatin A) overnight at room temperature. The pellet was then vortexed vigorously to dissolve. Each sample was vortexed for the same time and intensity. The solution was checked every minute by stereoscopic microscope to determine if the pellet had fully dissolved. A cutoff of 10 cycles of 1-minute vortexing was applied to define empirically the maximum amount of vortexing. The sample was centrifuged at room temperature at 20,000×g for 15 minutes to pellet detergent-insoluble proteins. The supernatant was collected as the 1% SDS or 1% sarkosyl soluble fraction. The remaining insoluble pellet was resuspended in 1 mL of 1% SDS or 1% sarkosyl lysis buffer and solubilized by vigorous vortexing using the same cutoffs. This was repeated until the third detergent-soluble fraction had been collected. The remaining pellet we define as the 1% SDS or 1% sarkosyl insoluble protein fraction. To redissolve the final pellet, 60 µL of 70% (v/v) formic acid was added before sonication in a Bioruptor at maximum intensity without cooling for 10 cycles of 1 minute on and 30 seconds off. This was followed by probe sonication at 70% intensity for 2 cycles of 20 seconds. The sample was then dried using a SpeedVac at maximum spin speed at 45°C. The resulting dried pellet was resolubilized in 2× LDS gel loading buffer with 5% (v/v) beta-mercaptoethanol and sonication in the Bioruptor for 5 cycles of 30 seconds on and 30 seconds off at maximum intensity. Samples were then boiled at 90°C for 10 minutes before storage at -20°C. The sample containing an unknown quantity of insoluble protein was then processed for mass spectrometry. An aliquot of the lysate (input) equivalent to 300 µg was processed by mass spectrometry in parallel.

## Mass Spectrometry Methods

### Proteolytic Digestion

The tissue homogenate or insoluble fraction was resuspended in 100 µL of 4% SDS in 50 mM triethylammonium bicarbonate buffer (TEAB) through vigorous mixing and incubated at 90°C for 1 hour. Samples were treated with 20 mM dithiothreitol in 50 mM TEAB at pH 7 at 56°C for 10 minutes, cooled to room temperature for an additional 10 minutes, and alkylated with 40 mM iodoacetamide in 50 mM TEAB (pH 7) at room temperature for 30 minutes in the dark. The samples were acidified by adding 6.5 µL of 12% phosphoric acid, and a fine precipitate formed after addition of 500 µL S-trap buffer and mixing by inversion. The entire mixture was added to S-trap spin columns (Protifi) and centrifuged at 4,000×g for 20 seconds or until fully eluted, followed by a wash with 400 µL of S-trap buffer. Samples were incubated with sequencing-grade trypsin (Promega) dissolved in 50 mM TEAB (pH 7) at a 1:25 enzyme:protein ratio for 1 hour at 47°C; additional trypsin was added at the same ratio and digestion proceeded at 37°C overnight. Peptides were eluted from the S-trap columns and dried by centrifugal evaporation, then resuspended in 800 µL of 1% formic acid in water and desalted using 10 mg Oasis SPE cartridges (Waters) for the lysate or ZipTips for the insoluble fractions. Desalted elutions were dried by centrifugal evaporation and resuspended in 20 µL of 0.2% formic acid in water for the insoluble fractions and in 100 µL of 0.2% formic acid in water for the lysate. Indexed Retention Time Standards (iRT, Biognosys) were added to each sample according to the manufacturer’s instructions.

### Chromatographic Separation and Mass Spectrometric Analysis

SDS-insoluble fractions and lysates were acquired on a Waters M-Class HPLC coupled to a ZenoTOF 7600 mass spectrometer (SCIEX) with an OptiFlow Turbo V Ion Source. The solvent system consisted of 0.1% formic acid in water (solvent A) and 99.9% acetonitrile, 0.1% formic acid in water (solvent B). Digested peptides were loaded onto a Luna Micro C18 trap column (20 × 0.30 mm, 5 µm particle size; Phenomenex) over 2 minutes at 10 µL/min using 100% solvent A, and eluted onto a Kinetex XB-C18 analytical column (150 × 0.30 mm, 2.6 µm particle size; Phenomenex) at 5 µL/min. Insoluble fractions were separated using a 45-minute linear gradient from 5 to 32% B, followed by an increase to 80% B for 1 minute, a hold at 80% B for 2 minutes, a decrease to 5% B for 1 minute, and a hold at 5% B for 6 minutes (55 minutes total acquisition time). Lysates were separated using a 120-minute linear gradient from 5 to 32% B with the same wash and re-equilibration steps (130 minutes total). For both, 4 µL of digested peptides were loaded. Ion source parameters were: gas 1 at 10 psi, gas 2 at 25 psi, curtain gas at 30 psi, CAD gas at 7 psi, source temperature 200°C, column temperature 30°C, positive polarity, spray voltage 5000 V. Data-independent acquisition (SWATH-DIA) was performed with survey MS1 spectra acquired over 395-1005 Da (accumulation time 100 ms, declustering potential 80 V, collision energy 10 V), with time bins to sum set to 8 and all channels enabled. MS2 spectra were collected over the same range using 80 variable windows (accumulation time 25 ms, dynamic collision energy, charge state 2, Zeno pulsing enabled). Total cycle time was 2.5 seconds.

Sarkosyl-insoluble fractions were acquired on a Dionex UltiMate 3000 system coupled to an Orbitrap Exploris 480 mass spectrometer (Thermo Fisher Scientific). The solvent system consisted of 2% acetonitrile, 0.1% formic acid in water (solvent A) and 80% acetonitrile, 0.1% formic acid (solvent B). Digested peptides (1:5 dilution) were loaded onto an Acclaim PepMap 100 C18 trap column (0.1 × 20 mm, 5 µm particle size; Thermo Fisher Scientific) over 5 minutes at 5 µL/min with 100% solvent A, and eluted onto an Acclaim PepMap 100 C18 analytical column (75 µm × 50 cm, 3 µm particle size; Thermo Fisher Scientific) at 300 nL/min using the following gradient: 2.5 to 24.5% B over 125 minutes, 24.5 to 39.2% B over 40 minutes, up to 98% B in 1 minute, and back to 2.5% B in 1 minute, with 30 minutes re-equilibration (210 minutes total). Each sample was acquired in data-independent acquisition (DIA) mode^78–80^. Full MS spectra were collected at 120,000 resolution (AGC target 3 × 10⁶ ions, maximum injection time 60 ms, 350-1,650 m/z) and MS2 spectra at 30,000 resolution (AGC target 3 × 10⁶ ions, automatic injection time, NCE 30, fixed first mass 200 m/z). The DIA isolation scheme consisted of 26 variable windows covering 350-1,650 m/z with 1 m/z overlap^79^.

### DIA-MS Data Processing, Quantification, and Statistical Analysis

Because SDS- and sarkosyl-insoluble fractions were acquired on independent instruments with distinct DIA schemes, all quantitative analyses were performed within acquisition runs; age-dependent fold changes were calculated independently for each detergent condition and were not compared in absolute magnitude across fractions.

SDS-insoluble and lysate data were analyzed using Spectronaut (version 17.6.230428.55965; Biognosys). The lysate was searched against the *H. sapiens* panhuman spectral library (PMID: 25977788, UniProt) with 20,526 entries (accessed 08/28/2015); the SDS-insoluble fraction was searched using the DirectDIA algorithm against the Human Reference Proteome (reviewed proteins only, UniProt) with 20,388 entries (accessed 07/31/2021). Sarkosyl-insoluble data were analyzed using Spectronaut (version 19.4.241104.62635; Biognosys) with DirectDIA against the *Homo sapiens* reference proteome with 20,411 entries (UniProtKB-SwissProt, accessed 10/06/2023).

For all searches, dynamic data extraction parameters and precision iRT calibration with local (SDS) or non-linear (sarkosyl) regression were used. Trypsin/P was specified as the digestion enzyme, allowing for specific cleavages and up to two missed cleavages. Methionine oxidation and protein N-terminus acetylation were set as dynamic modifications, while carbamidomethylation of cysteine was set as a static modification. Protein identification at the group level required at least 2 unique peptide identifications and was performed using a 1% q-value cutoff for both precursor ion and protein levels. Quantification was based on the peak areas of extracted ion chromatograms (XICs) of 3-6 MS2 fragment ions, specifically b- and y-ions, with 1% q-value data filtering applied. For lysate data, cross-run normalization (SDS acquisition) or local normalization (sarkosyl acquisition) was applied; insoluble fraction data were not normalized, to avoid bias driven by differences in total protein quantity across samples. Relative protein abundance changes were compared using paired t-tests with p-values corrected for multiple testing using the Storey method^81^ with group-wise testing corrections. Proteins with at least two unique peptides, q < 0.05, and |log₂(fold change)| > 0.58 were considered significantly altered.

### Western Blotting

To validate mass spectrometry findings, western blotting was performed on SDS-insoluble and sarcosyl-insoluble pellet fractions from a subset of the same cohort used for proteomic analysis. Pellet fractions were resuspended in 2xLDS buffer and boiled for 10 minutes, before equal volumes were loaded onto 4-12% Bis-Tris gels and resolved by SDS-PAGE. Proteins were then transferred to nitrocellulose membranes via semi-dry transfer using a Biorad transblot turbo. Membranes were blocked in 5% non-fat milk in TBS-T for 1 hour at room temperature, then incubated overnight at 4°C on a rocker with primary antibodies against PIN1 (mouse monoclonal, 1:2000, Cat. No. 68127-1-Ig, Proteintech), Cystatin C (CST3 rabbit polyclonal, 1:6000, Cat. No. 12245-1-AP, Proteintech), total tau (Tau5, mouse monoclonal, 1:500, Cat. No. AHB0042, Invitrogen), and HTT (rabbit polyclonal, N-terminal, 1:150, Cat. No. H7540, Sigma-Aldrich). Following primary antibody incubation, membranes were washed in TBS-T and incubated with HRP-conjugated secondary antibodies (anti-mouse or anti-rabbit as appropriate; 1:2000 dilution) for 2 hours at room temperature. Protein bands were visualised using Thermo Scientific SuperSignal West Pico kit and imaged on a Bio-Rad ChemiDoc system. Band intensities were quantified by densitometry using FIJI (ImageJ). As equal volumes rather than equal protein concentrations were loaded per fraction, band intensities were not normalized to a loading control; this approach preserved the native protein yield differences between samples, consistent with the approach applied to the mass spectrometry data.

## Computational Methods

### Functional Enrichment Analysis

All functional enrichment analysis was performed using the gProfiler web server^82^. For GO and pathway enrichment analyses, the applicable experimentally measured background universe was incorporated, and the Benjamini-Hochberg method was used with an FDR correction threshold of 5%.

### Monotonic Trend Test on Sarkosyl-insoluble Accumulators

Monotonic trend was tested per protein using MannKendall() from the Kendall R package on abundance values ordered by continuous age; p-values were adjusted for multiple testing using the Benjamini-Hochberg procedure across the 303 sarkosyl-insoluble accumulators.

### Protein Trajectory Modeling and Clustering

To characterize age-related changes in protein abundance and insolubility, we fitted three regression models to each protein: linear regression (abundance ∼ age), natural cubic spline regression (df = 3), and generalized additive models (GAM with penalized thin-plate spline, k = 10-15, REML optimization). For each protein, abundance values were first z-scored within protein to enable cross-protein comparison.

Model performance was assessed using adjusted R² (linear, spline) or deviance explained (GAM), root mean square error (RMSE), and smooth term p-value. The best-fitting model was selected as that with highest adjusted R²/deviance explained. Technical variability was quantified as the coefficient of variation (CV) of raw abundance across individuals. Proteins were filtered for trajectory clustering based on quantile thresholds applied to model fit metrics: adjusted R² > 75th percentile, RMSE < 75th percentile, CV > 25th percentile (to retain dynamically changing proteins), and p < 0.05. These thresholds were calculated separately for each fraction (lysate, SDS-insoluble, sarkosyl-insoluble) and for aggregation ratio (insoluble/lysate) analyses. Filtered proteins were clustered by k-means (k = 6, seed = 42) based on mean z-scored values within discrete age groups: Young (20-35 yrs), Middle (45-55 yrs), Old (65-75 yrs), and Geriatric (80-88 yrs).

### Robustness Estimations

Leave-one-out (LOO) analysis was performed to assess sensitivity of age-related changes to individual subjects. For each fraction, we iteratively excluded each geriatric subject (n = 14 for SDS, n = 11 for sarkosyl) and recalculated the median log2 fold change (Geriatric vs Young) across all detected proteins. Robustness was evaluated using sign consistency, signal-to-noise ratio, coefficient of variation, and 95% confidence intervals derived from the LOO distribution. To test whether influential subjects showed evidence of preclinical AD pathology, we calculated Pearson correlations between LOO influence and sarkosyl-insoluble abundance of APP and MAPT.

Because biobank source and age group were partially confounded in this cohort, with younger subjects drawn predominantly from one consortium site and older subjects from another, we did not include biobank as a covariate. Post-mortem interval, the primary technical variable associated with tissue banking, was modeled directly. We fitted per-protein linear models (log2 abundance ∼ age + PMI), with age as a continuous variable, for all proteins reaching significance in the primary analysis (q < 0.05, |log2FC| > 0.58, Geriatric vs Young) and tested whether the PMI coefficient reached nominal significance. PMI reached nominal significance (p < 0.05) for 5.0% and 0.6% of SDS and sarkosyl proteins respectively. Within age groups represented by multiple biobank sites (minimum 2 subjects per site), abundance profiles were highly correlated across sites for both SDS-insoluble and sarkosyl-insoluble fractions, and for both the full detected proteome and age-changing proteins (Pearson r = 0.931-0.979 across 30 pairwise comparisons), indicating that site of origin does not introduce systematic bias. We further tested whether age effects were reproducible within individual biobank sites by comparing mean log2 abundance between adjacent age groups within Maryland (Young vs Middle) and Harvard (Old vs Geriatric); 98.0% of significant sarkosyl-accumulating proteins (297 of 303) increased from Old to Geriatric within Harvard subjects, and 95.0% (288 of 303) increased from Young to Middle within Maryland subjects.

### Cell Composition Analysis

Cell proportion estimation was performed using the BrainDeconvShiny App website, which uses the dTangle algorithm to estimate brain cell proportions based on published single cell/nuclei datasets^83^. We used the consensus reference panel ’MultiBrain,’ which brings together five independent studies characterizing brain cell expression profiles, to estimate cell proportions based on lysate protein abundance data^83^. dTangle achieved a goodness of fit of approximately 0.4.

### Biophysical Feature Enrichment Analysis

Proteins were annotated with biophysical and post-translational modification features. Continuous features included: intrinsic disorder content, classical aggregation propensity (TANGO score, Zagg ^42,43^), supersaturation (taken from^45^), LLPS propensity (catGRANULE, FuzDrop ^46,47^), and predicted solubility (CamSol variant score^44^). Sequence length, isoelectric point (pI), hydrophobicity (GRAVY), and secondary structure propensities (helix, sheet, turn) were taken from UniProt^48^. Categorical features included: RNA-binding capacity (Gene Ontology), chaperone-mediated autophagy (CMA) targeting motif presence ^50^, ubiquitination site type (K48, K63), phosphorylation, and glycosylation status were taken from Uniprot^48^.

For intrinsic disorder content, per residue disorder assignments for the human reference proteome were obtained from AlphaFold2^49^ predictions and collapsed into contiguous disordered regions, each represented as a start–end residue range. For every protein we computed the fraction of residues falling within disordered regions (percent disorder) by summing the lengths of all annotated regions and dividing by total protein length; proteins without any predicted disordered region were assigned a value of zero.

For continuous features, group distributions were compared using Wilcoxon rank-sum tests. Categorical features were compared using Fisher’s exact tests. For each feature, z-scores were calculated to standardize effect sizes across continuous and categorical feature types. Therefore, for continuous features, the z-score was defined as the difference between the group mean and the reference mean, divided by the reference standard deviation, and for categorical features, the z-score was defined as the difference in proportions, standardized by the expected binomial variation of the reference proportion. P-values were corrected for multiple testing within each comparison group using Benjamini-Hochberg FDR correction

### Multivariate analysis of biophysical predictors

To test whether LLPS predictor enrichment among sarkosyl-accumulating proteins could be explained by correlated features such as sequence length, we performed logistic regression with accumulator status as the outcome variable. The background population consisted of all proteins detected in the sarkosyl-insoluble fraction (n = 2,208 with complete feature annotations). Accumulators were defined as proteins showing significant increase in the Geriatric versus Young comparison (q < 0.05, log₂ fold change > 0.58; n = 238 in the model). We first fitted a detection-only model including sequence length, lysate abundance, and unique peptide counts as predictors. We then fitted a full model adding LLPS propensity scores (catGRANULE, FuzDrop). All continuous predictors were z-scored prior to model fitting. Sequence length, which showed marginal association with accumulation in the detection-only model (p = 0.074), was no longer significant after including LLPS predictors (p = 0.74). Both catGRANULE and FuzDrop remained significant after controlling for sequence length, abundance, and peptide counts (catGRANULE β = 0.17, p = 0.042; FuzDrop β = 0.26, p = 4.9 × 10⁻⁴).

### Alzheimer’s Disease Pathology Enrichment

AD-associated proteins were obtained from the NeuroPro database, which has plaqueome and tangleome protein sets^56^. CSF biomarkers were proteins taken from a meta-analysis of CSF biomarkers and filtered to those from at least three independent studies^57^. Overlap between each insoluble proteome and the AD proteomes was quantified using Fisher’s exact tests, with fold enrichment calculated relative to the hippocampal proteome to visualize baseline enrichment and aging-specific incremental enrichment. All p values were adjusted using the Benjamini-Hochberg FDR method. Venn/Euler diagrams were generated using the eulerr package in R to visualize set relationships.

### Enrichment of proteostasis network and cell type-specific proteins in the insoluble proteome

We used Fisher’s exact tests to assess whether proteins from specific proteostasis network (PN) branches or cell types were over-represented among (i) proteins detected in each insoluble fraction relative to the lysate-detected proteome, and (ii) age-changing proteins within each fraction relative to all detected proteins in that fraction. PN annotations were obtained from the Proteostasis Network Consortium database^61,62^ and restricted to proteins with exclusive single-branch membership across nine branches (mitochondrial, autophagy, translation, ER, cytosolic chaperones, lysosomal, proteasome/UPS, extracellular, nuclear). Cell type markers for six brain cell types were obtained from the BRETIGEA R package^63^.

### Proteostasis network and cell type predictors of insoluble protein burden

Two per-individual outcome variables were defined: summed log10-transformed abundance of age-decreasing SDS-insoluble proteins and age-increasing sarkosyl-insoluble proteins (q < 0.05, |log2FC| > 0.58, Geriatric vs Young). For each PN branch and cell type, a per-individual signature score was calculated as the mean z-scored lysate abundance of all detected branch members. Each score was tested in a linear model adjusting for age. BRETIGEA cell type signatures were tested using the same framework. To distinguish mitochondrial proteostasis capacity from overall mitochondrial abundance, total mitochondrial protein content was estimated using the annotations from MitoCarta 3.0^84^ and tested as an alternative predictor.

The translation branch was decomposed into eight mutually exclusive subcategories by function and compartment: cytosolic and mitochondrial ribosomal proteins, cytosolic initiation factors, cytosolic elongation/termination factors, cytosolic aminoacyl-tRNA synthetases, mitochondrial translation factors, and cytosolic and mitochondrial ribosome biogenesis factors. Additionally, six co-translational proteostasis categories were defined: nascent chain chaperones (NAC/RAC), ribosome quality control (RQC), UFMylation, mitochondrial import (TOM/TIM), ER import (SRP/SEC61), and the TRiC/CCT chaperonin. TR breakdown was tested independently of the nine-branch framework. Per-individual signature scores were calculated as the mean z-scored lysate abundance of all detected subcategory members.

To assess multicollinearity among proteostasis predictors in joint models, we calculated variance inflation factors (VIF) using the car package in R; all VIF values were below 3, well under the threshold of 5-10 typically considered problematic for coefficient stability.

## Data Availability

Raw proteomics data and complete MS data sets will be uploaded to the Mass Spectrometry Interactive Virtual Environment (MassIVE) repository, developed by the Center for Computational Mass Spectrometry at the University of California San Diego, and can be downloaded using the following link: (MassIVE ID number: ProteomeXchange ID:)

## Code Availability

Analysis scripts used to generate figures, tables, and statistical results reported in this manuscript, together with the source code for the accompanying Shiny application, are hosted at https://github.com/Eanderton91/human_hippocampus_insoluble_proteome and archived at Zenodo (DOI: 10.5281/zenodo.19617066.) under an MIT license. The Human Hippocampus Insoluble Proteome Explorer, an interactive web application for browsing per-protein age trajectories, biophysical features, and AD pathology overlap, as well as downloading the complete human proteome with biophysical feature assignments is hosted at https://jzezxu-edward-anderton.shinyapps.io/human-hippocampus-insoluble-proteome-explorer/. All scripts were written in R (version 4.4.1).

## Acknowledgements

We thank the NIH NeuroBioBank for providing post-mortem brain samples. This research was supported by NIH grants RFAG057358 (G.J.L), and R01AG02963 (G.J.L), the Larry L. Hillblom Foundation LLHF Ctr Supp. and NIA U01AG045844 (G.J.L), the University of Southern California and Buck Institute Nathan Shock Center P30AG068345 (MPI: Verdin, Curran, Cohen, Lithgow) and Cellular Senescence and Beyond Core (Schilling) (E.A), and The Diana Jacob Kalman American Federation for Aging Research (AFAR) Scholarship (E.A).

## Ethics Statement

Collection and distribution procedures at each Brain and Tissue Repository are conducted under IRB-approved protocols at their respective institutions. Because this study used de-identified post-mortem tissue, it was determined not to constitute human subjects research and was exempt from additional IRB review.

---

**Supplementary Figure 1.**
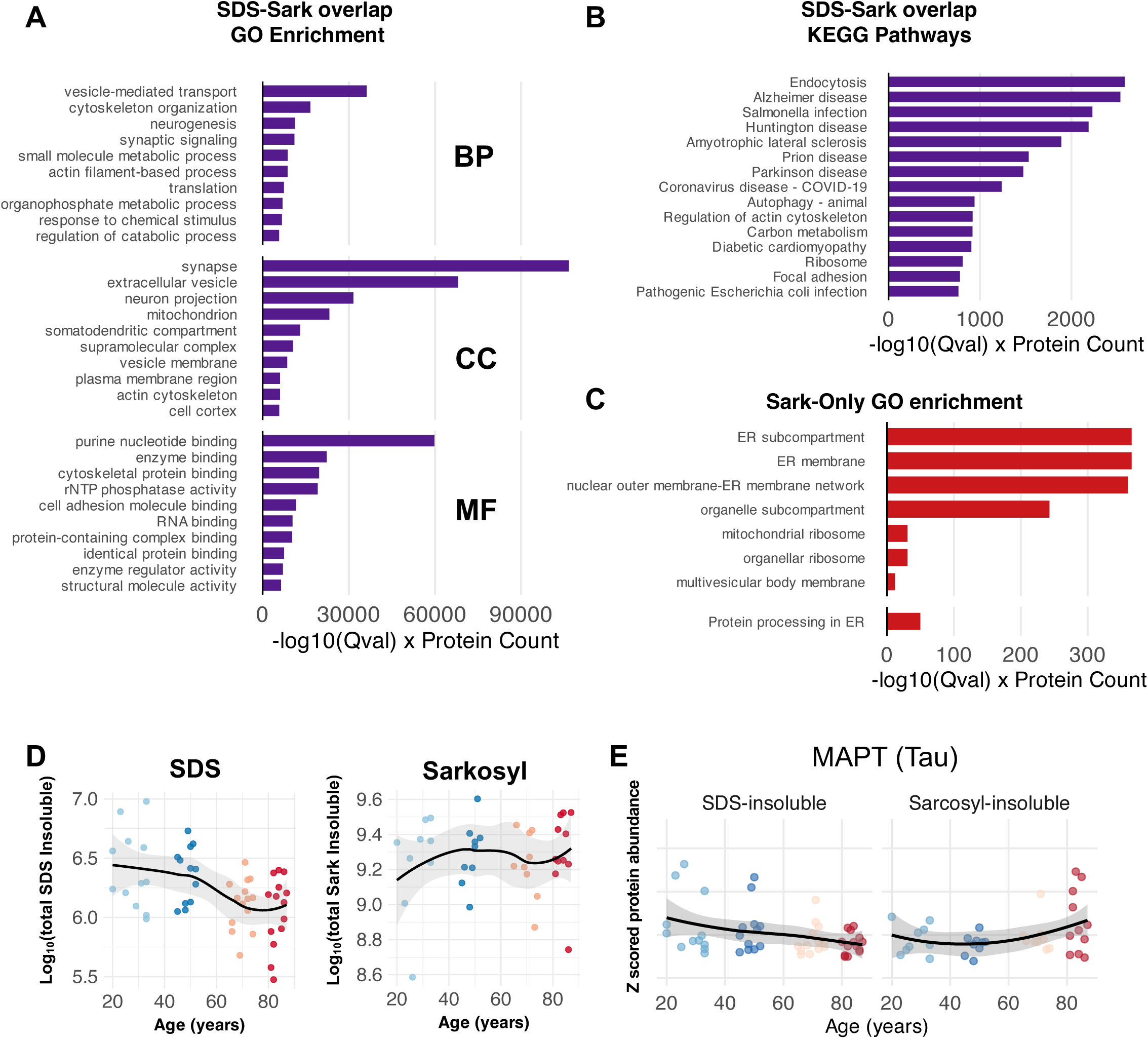
**A.** Barplot of top enriched GO terms for insoluble proteins common to both SDS and Sarkosyl fractions compared against the hippocampal proteome background, all q<0.05, Fisher’s one-tailed test with Benjamini-Hochberg FDR correction. **B** As in (A) with top enriched KEGG pathway terms. **C.** Barplot of top enriched GO terms for insoluble proteins measured exclusively in the Sarkosyl fraction compared against the Sarkosyl insoluble detected as background, all q<0.05, Fisher’s one-tailed test with Benjamini-Hochberg FDR correction. **D.** Loess plots of Log_10_(Total Insoluble Signal) against subject age for SDS insoluble proteome (left) and Sarkosyl insoluble proteome (right), each data point represents an independent biological sample (individual donor). **E.** Loess plots of z-scored insoluble Tau protein abundance against subject age for SDS (left) and Sarkosyl (right), each data point represents an independent biological sample (individual donor).

**Supplementary Figure 2.**
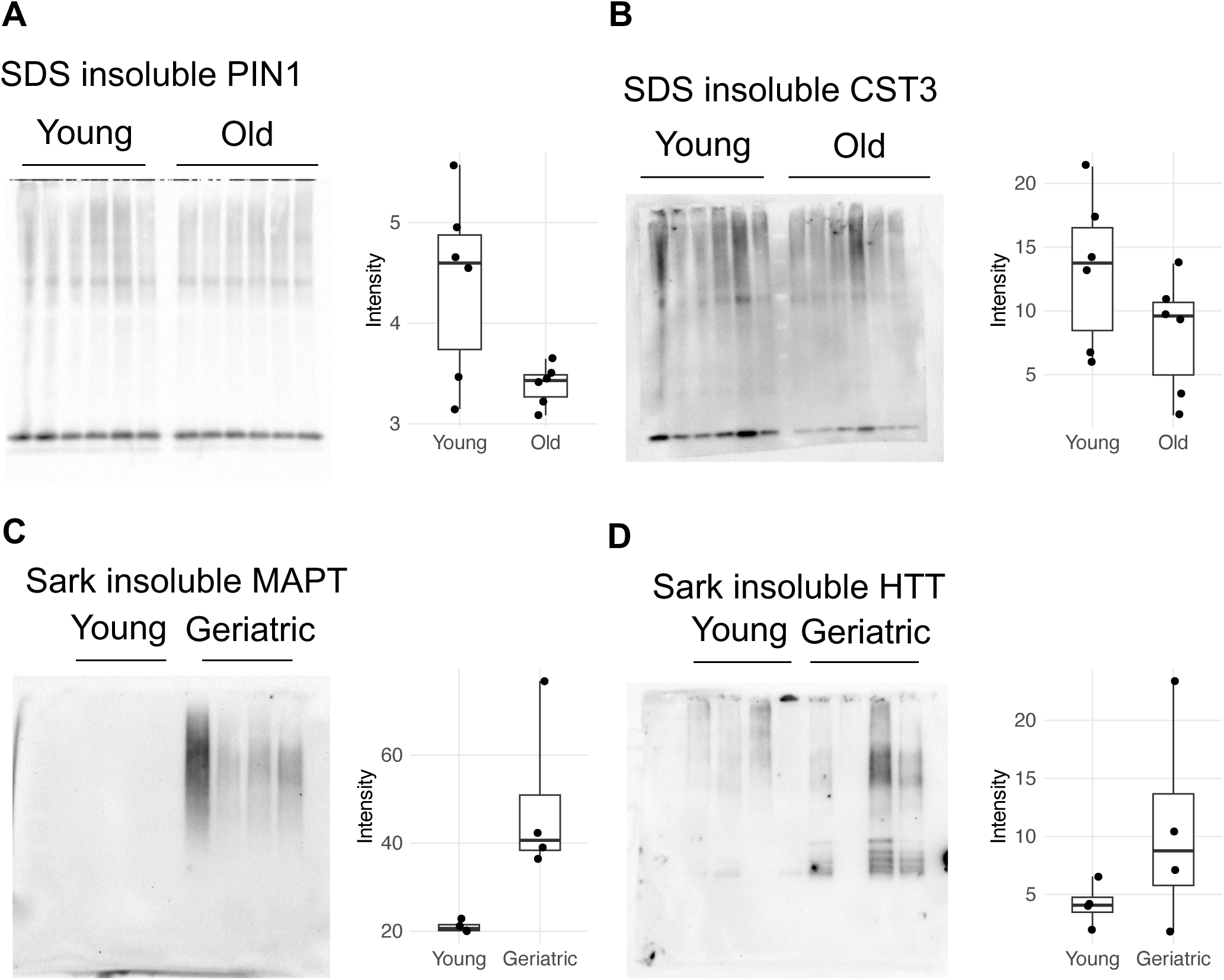
**A**. Western blot of SDS insoluble PIN1, 6 young and 6 old. **B**. Western blot of SDS insoluble CST3, 6 young and 6 old. **C**. Western blot of Sarkosyl insoluble MAPT (Tau), 4 young and 4 geriatric. **D**. Western blot of sarkosyl insoluble HTT, 4 young and 4 geriatric. For all blots each lane represents an independent biological sample (individual donor).

**Supplementary Figure 3.**
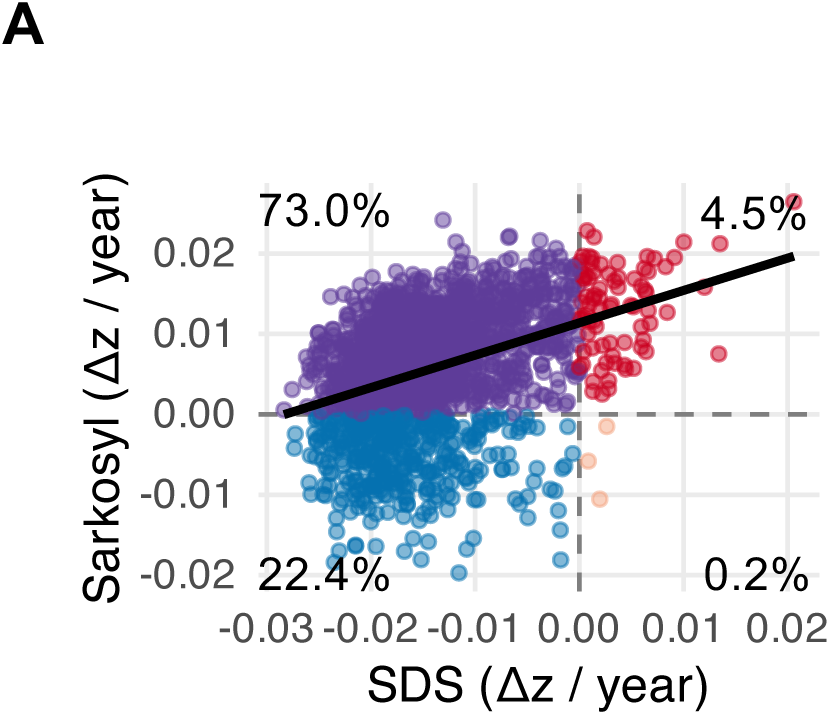
A Scatter plot for all proteins detected in both fractions showing Δz-score mean change over aging in the Sarkosyl fraction and SDS fraction. Spearman correlation line shows positive correlation.

**Supplementary Figure 4.**
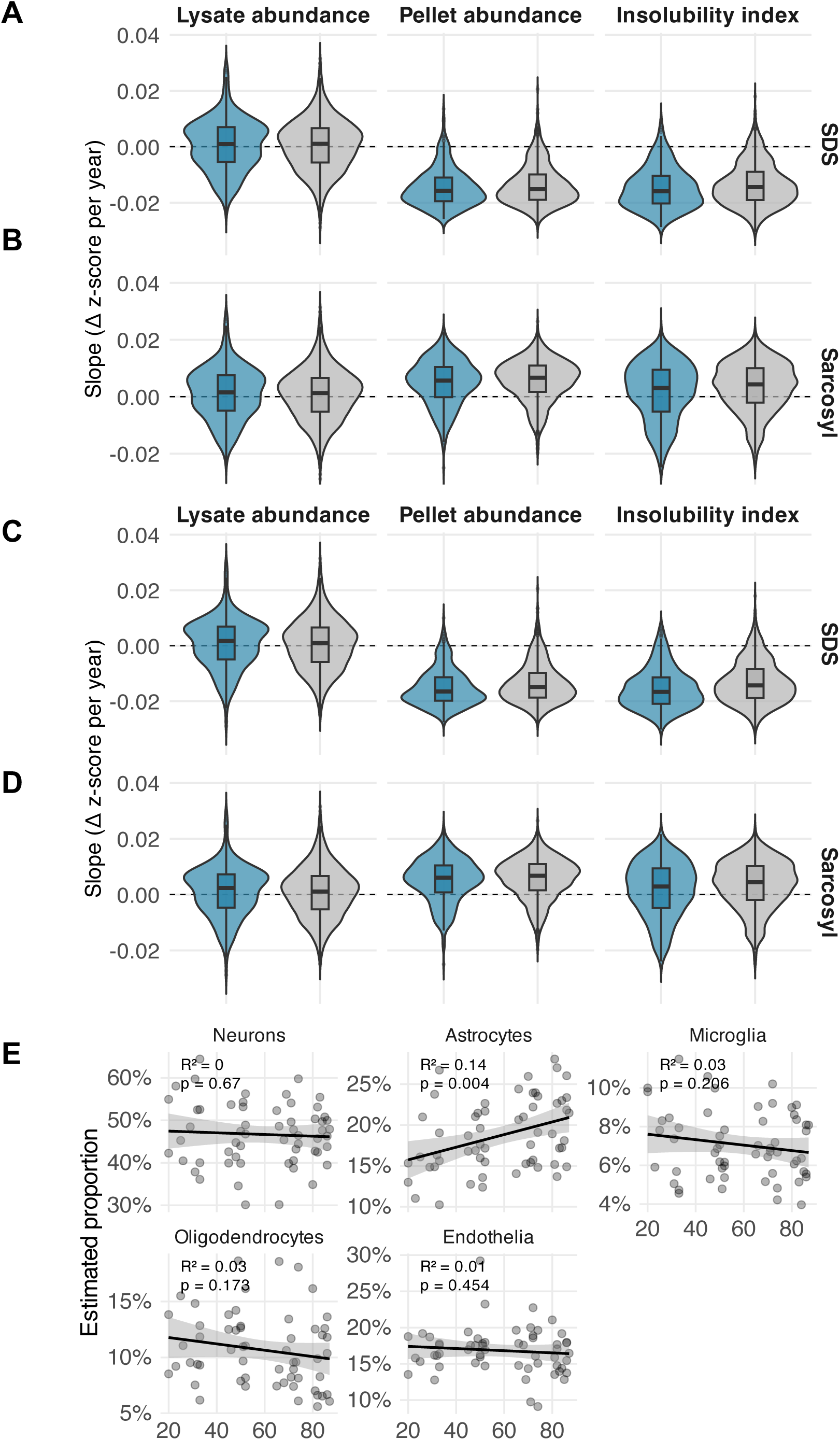
**A** Violin plot showing Δz-score mean change over aging for synaptic SDS insoluble proteins vs SDS insoluble detected proteins in terms of abundance in the pellet, abundance in the lysate, and the insolubility index (pellet / lysate), Mann Whitney U test. **B**. As in (A) with synaptic Sarkosyl insoluble proteins. **C-D**. As in (A-B) with post-synaptic density (PSD) insoluble proteins, Mann Whitney U test. **E.** Cell type proportion estimates for major central nervous system cell types. Pearson correlation analysis, estimated using dTangle and multibrain human reference signature, each data point represents an independent biological sample (individual donor). **A** Overlap between detected insoluble proteomes and plaquome. **B** Overlap between detected insoluble proteomes and tangleome. **C**. Overlap between age-changing proteins and plaqueome for SDS insoluble proteins (left) and Sarkosyl insoluble proteins (right). **D** As in © with tangleome for both fractions. Yellow = plaques, Green = Tangles, Blue = SDS, Red = Sarkosyl.

**Supplementary Figure 5.**
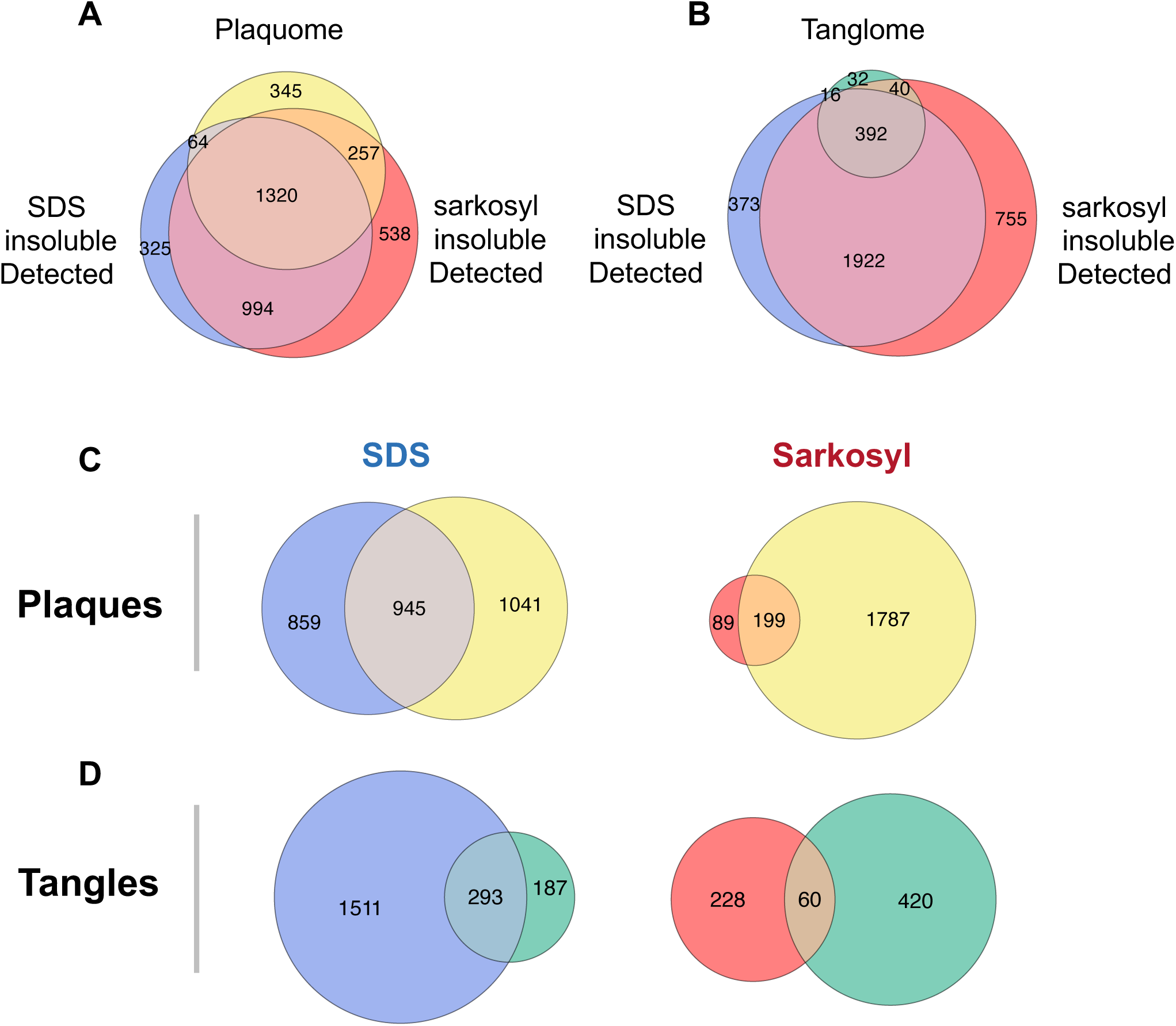
**A** Overlap between detected insoluble proteomes and plaquome. **B** Overlap between detected insoluble proteomes and tangleome. **C**. Overlap between age-changing proteins and plaqueome for SDS insoluble proteins (left) and Sarkosyl insoluble proteins (right). **D** As in © with tangleome for both fractions. Yellow = plaques, Green = Tangles, Blue = SDS, Red = Sarkosyl.

**Supplementary Figure 6.**
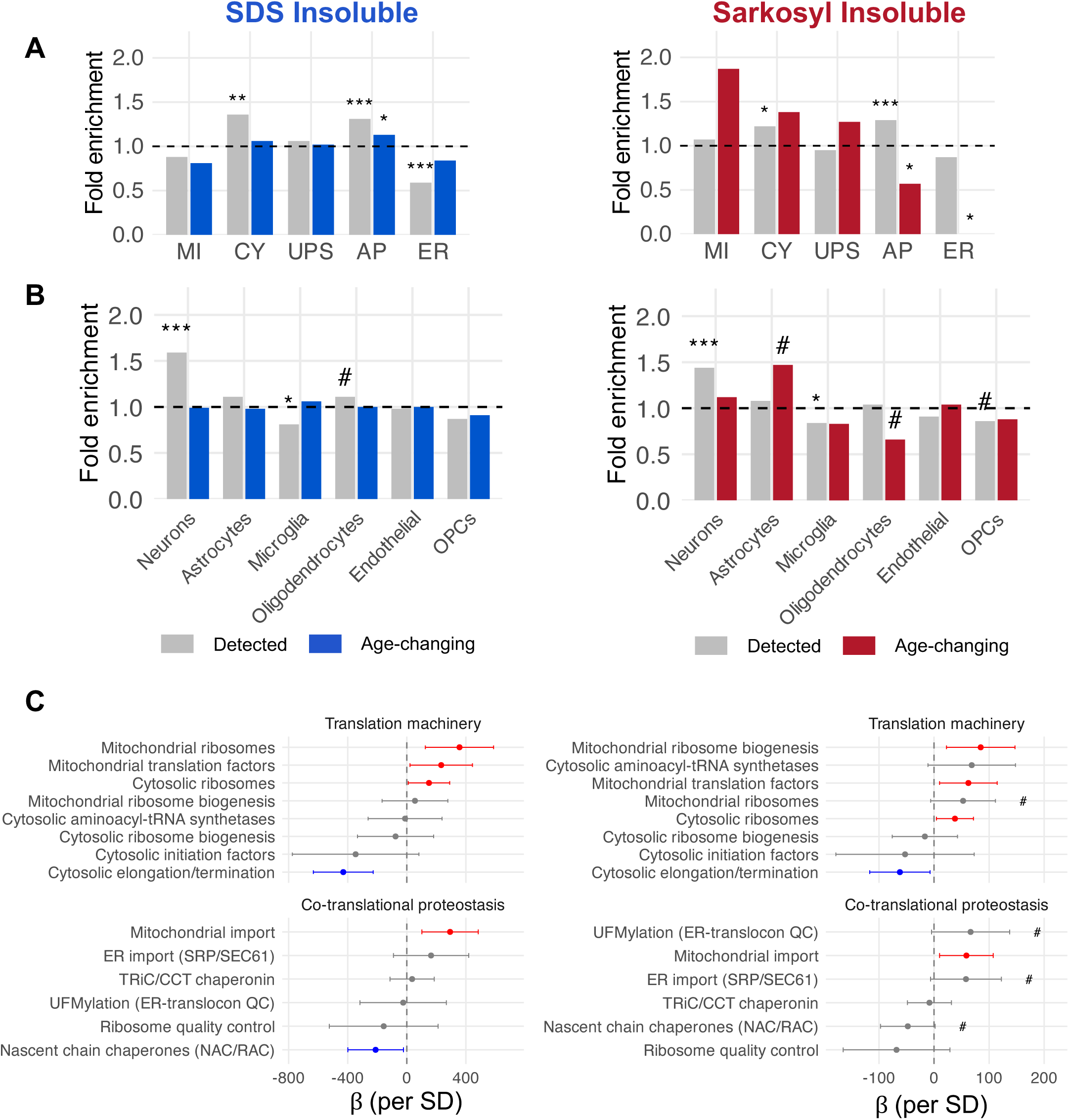
**A** Fold enrichment of proteostasis branch proteins in the insoluble proteome, Left = SDS, Right = Sarkosyl (Grey bars: insoluble detected, Blue: SDS age-changing, Red: Sarkosyl age-changing. **B** As in A, fold enrichment but using cell type signature scores and insoluble burden. Fisher’s exact test with Benjamini-Hochberg correction. **C.** Forest plots showing β coefficient of age-adjusted linear models for translation-associated proteostasis network branch broken down into sub-branches left = SDS, right = Sarkosyl. Points show the beta coefficient +/- 95% confidence interval from a linear model adjusting for age (insol. burden ∼ age + predictor). Red: significant positive (p < 0.05); Blue: significant negative (p < 0.05); grey, non-significant. # = p<0.1. MI = mitochondrial, CY = cytoplasmic, UPS = proteasomal, AP = autophagy. ***q < 0.001, **q < 0.01, *q < 0.05, #q <0.1. QC = Quality Control.

